# Lysosomal integrity suppresses TIR-1/SARM1 aggregation to restrain toxic propagation of p38 innate immunity

**DOI:** 10.1101/2024.02.16.580734

**Authors:** Samantha Y. Tse-Kang, Read Pukkila-Worley

## Abstract

Innate immunity in bacteria, plants and animals requires the specialized subset of TIR-domain proteins that are NAD^+^ hydrolases. Aggregation of these TIR proteins engages their enzymatic activity, but it is not known how this protein multimerization is regulated. Here, we discovered that TIR oligomerization is exquisitely controlled to prevent immune toxicity. We found that p38 propagates its own activation by promoting the feedforward expression and aggregation of the lone enzymatic TIR protein in the nematode *C. elegans*, TIR-1/SARM1. We performed a forward genetic screen to determine how the p38 positive feedforward loop is regulated. We discovered that the integrity of the specific lysosomal sub-compartment that expresses TIR-1/SARM1 is actively maintained to limit inappropriate aggregation of this protein and restrain toxic p38 immune activation. Thus, innate immune defenses in intestinal epithelial cells are regulated by specific control of TIR-1/SARM1 multimerization.

## INTRODUCTION

Tight control of inflammation is essential for animal health and is a particular challenge for cells in barrier tissues, such as the intestinal epithelium, that constantly interface with both commensal organisms and virulent pathogens. Conceptually, pathogen sensing mechanisms, which activate protective inflammatory signaling cascades during infection, must themselves be regulated to prevent pathology associated with exuberant immune activation. How this occurs, however, is not fully understood.

In this study, we define an immunoregulatory axis anchored by a protein whose function in innate immunity is conserved across the tree of life. A specific subset of proteins that contain toll/interleukin-1/resistance gene (TIR) domains are enzymes that metabolize nicotinamide adenine dinucleotide (NAD+) to produce secondary metabolites.^1–3^ Like other proteins with a TIR domain, which include Toll-like receptors (TLRs) in animals, these proteins are essential for coordinating host defense against pathogen infection. The catalytic activity of plant TIR, for example, is required to induce protective cell death responses during pathogen infection.^2^ Enzymatic TIR proteins also function in bacterial immunity, orchestrating protective anti-phage defenses.^4^ It is noteworthy that the enzymatic activity of TIR is often dependent on protein oligomerization. For example, plant TIR forms a tetrameric structure upon activation that drives its catalytic activity.^5,6^ In addition, prokaryotic TIR-containing proteins oligomerize into filaments to induce NAD+ hydrolase activity.^3^ A TIR-containing protein with enzymatic activity in animals and humans, sterile alpha and TIR motif-containing 1 (SARM1), requires multimerization to engage its intrinsic NAD+ hydrolase activity.^7,8^ Oligomerization of enzymatic TIR proteins is therefore conserved across millions of years of evolution; however, it is not known how TIR multimerization is regulated.

Toll/interleukin-1/resistance gene (TIR)-1 in the nematode *C. elegans* is the homolog of mammalian SARM1. TIR-1/SARM1 functions upstream of the p38 PMK-1 mitogen activated protein kinase (MAPK) and is required for host defense against infection with a variety of pathogens.^9–11^ Previously, we discovered that TIR-1/SARM1 multimerizes in intestinal epithelial cells during pathogen infection.^12^ Oligomerization of TIR-1/SARM1 in this manner engages its intrinsic NAD+ hydrolase activity^8,12,13^ to activate the p38 PMK-1 pathway.^12^ In a companion manuscript co-submitted with this study, we discovered that TIR-1/SARM1 is expressed on the membranes of a specific subset of lysosomes called lysosome-related organelles. Intracellular oxidative stress, induced by the redox-active bacterial virulence effector pyocyanin, collapsed lysosome-related organelles, which in turn led to aggregation and activation of TIR-1/SARM1. Pyocyanin induced TIR-1/SARM1 multimerization on the membranes of lysosome-related organelles, which activated p38 PMK-1 signaling. Thus, lysosomal TIR-1/SARM1 senses a pattern of pathogenesis to identify bacterial infection and activate innate immunity.

Here, we discovered that aberrant TIR-1/SARM1 aggregation and activation in the intestine is toxic and is therefore tightly controlled. Inappropriate activation of p38 PMK-1 is a well-known contributor to pathology in nematodes and mammals. We demonstrated that these immune toxicities are regulated at the level of TIR-1/SARM1. We further showed that lysosome-related organelle integrity is actively maintained to suppress TIR-1/SARM1 multimerization and restrain toxic p38 PMK-1 activation in *C. elegans*. We found that p38 PMK-1 drove a positive feedforward loop, which potentiated its own activation by promoting the expression, aggregation, and activation of TIR-1/SARM1. We performed a forward genetic screen to determine how feedforward activation of p38 PMK-1 is controlled and identified a previously uncharacterized gene, which we named regulator <u>o</u>f *<u>tir</u>-1* (*rotr-1)*. ROTR-1 maintained the size of lysosome-related organelles, which promoted immune homeostasis by limiting aggregation and activation of TIR-1/SARM1. Thus, our unbiased forward genetic screen identified *C. elegans* mutants that recapitulated the pathogen-induced damage to the specific lysosomal compartment that expresses TIR-1/SARM1, providing orthologous confirmation for the cell biological characterization of p38 PMK-1 immune activation in our companion manuscript. In addition, these data demonstrate that a lysosomal – TIR-1/SARM1 – p38 axis restrains immune activation in intestinal epithelial cells to promote healthy growth and longevity of *C. elegans*.

## RESULTS

### Feedforward activation of the p38 PMK-1 pathway potentiates innate immune defenses by promoting TIR-1/SARM1 multimerization

*C. elegans* TIR-1/SARM1 activates the p38 PMK-1 immune pathway, a classic MAPK signaling cassette with the MAPKKK NSY-1 (homolog of mammalian ASK1), the MAPKK SEK-1 (homolog of mammalian MKK3/6), and p38 PMK-1. The phosphatase *vhp-1* dephosphorylates p38 PMK-1 to negatively regulate innate immune defenses (**Figure S1A**).^14^ We examined the mechanisms of TIR-1/SARM1 regulation and made the surprising observation that RNAi-mediated knockdown of *vhp-1*, which hyperactivates p38 PMK-1,^14^ caused robust induction of TIR-1/SARM1 protein expression (**Figures 1A-B**). This result was unexpected because p38 PMK-1 is downstream of TIR-1/SARM1 in the activation of innate immune defenses (**Figure S1A**).^9^ For these studies, we used a *C. elegans* strain, previously engineered in our laboratory using clustered regularly interspaced short palindromic repeats (CRISPR)-Cas9, that expresses TIR-1 protein tagged from its endogenous locus with a 3xFLAG sequence (TIR-1::3xFLAG).^12^ These data suggest that p38 PMK-1 may potentiate its own activation in a positive feedforward cycle to propagate activation of anti-pathogen defenses.

**Figure 1.**
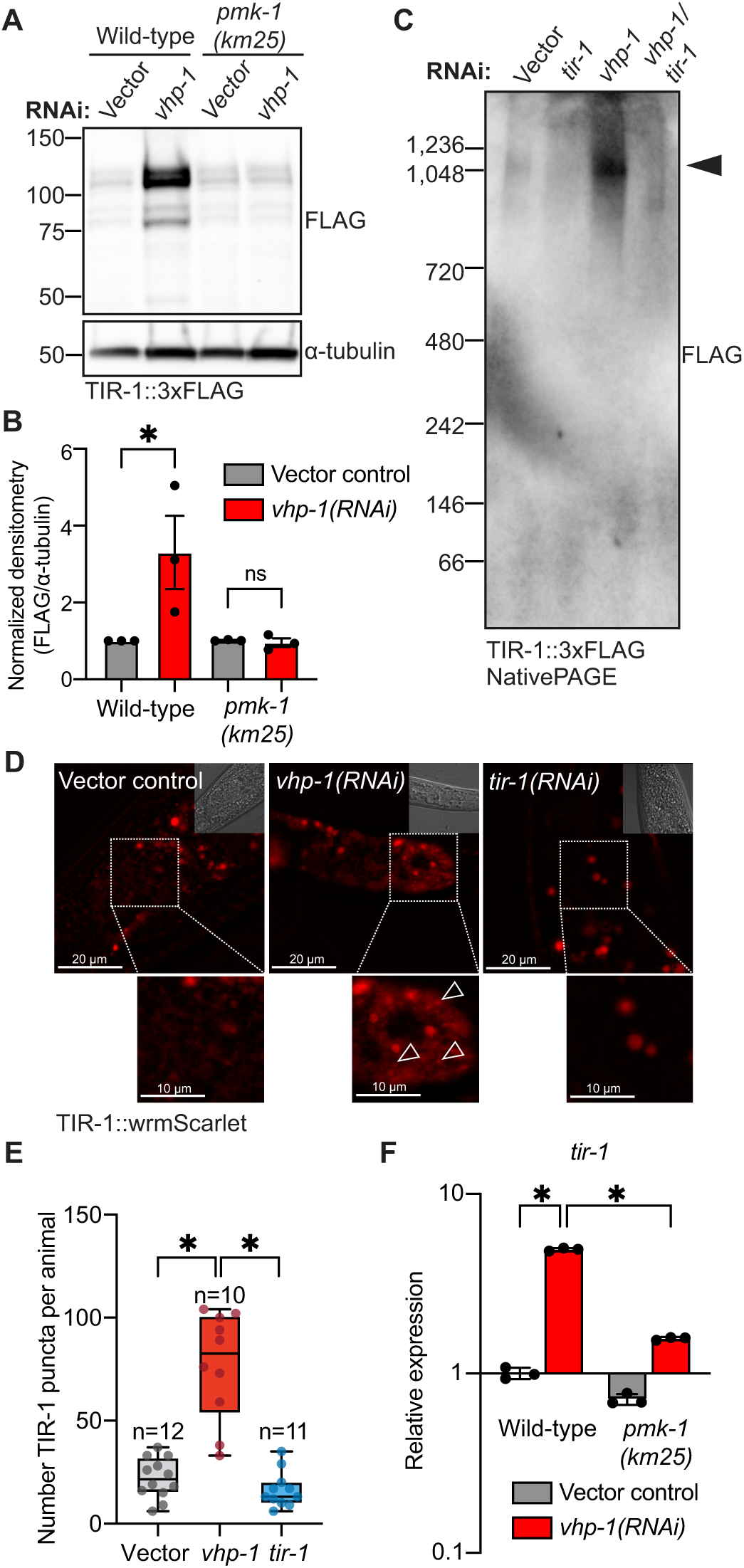
Feedforward activation of the p38 PMK-1 pathway potentiates innate immune defenses by promoting TIR-1/SARM1 multimerization. **(A)** Representative immunoblot using an anti-FLAG antibody on whole cell lysates of wild-type and *pmk-1(km25)* mutants in the TIR-1::3xFLAG background treated with vector control or *vhp-1(RNAi)*. **(B)** Densitometric quantification of (A). Error bars represent SEM (*n*=3). *equals p<0.05 (two-way ANOVA with Šídák’s multiple comparisons test). **(C)** Blue NativePage immunoblot on *C. elegans* TIR-1::3xFLAG animals treated with vector control, *tir-1(RNAi), vhp-1(RNAi),* or *vhp-1/tir-1* double RNAi. Immunoblot was probed using the anti-FLAG antibody. Arrow indicates TIR-1 multimer. **(D)** Images of *C. elegans* TIR-1::wrmScarlet animals treated with vector control, *vhp-1(RNAi)* or *tir-1(RNAi)*. Insets represent corresponding DIC images. Dotted boxes indicate higher magnifications. Open arrowheads indicate TIR-1 puncta. **(E)** Quantification of the number of TIR-1::wrmScarlet puncta in (D), as previously described.^12^ *equals p<0.05 (one-way ANOVA with Tukey’s multiple comparisons test). **(F)** qRT-PCR analysis of *tir-1* transcription in wild-type and *pmk-1(km25)* mutant animals treated with vector control or *vhp-1(RNAi).* Error bars represent SD. *equals p<0.05 (two-way ANOVA with Tukey’s multiple comparisons test). Source data for this figure is in Table S3. See also Fig. S1.

Activation of p38 PMK-1 requires multimerization of TIR-1/SARM1 into protein assemblies (puncta) in intestinal epithelial cells, which engages its intrinsic NAD^+^ hydrolase activity.^12^ We therefore explored whether *vhp-1(RNAi)* promoted TIR-1/SARM1 aggregation. First, we examined TIR-1::3xFLAG under non-denaturing, or native, conditions. RNAi of *vhp-1* increased TIR-1/SARM1 protein expression and induced the formation of higher-order multimers (**Figure 1C**). Second, we used a *C. elegans* strain that we previously engineered using CRISPR-Cas9, which expresses TIR-1/SARM1 protein tagged with the fluorophore wrmScarlet at its endogenous genomic locus.^12^ RNAi-mediated knockdown of *vhp-1* increased TIR-1::wrmScarlet protein expression and significantly induced the formation of TIR-1::wrmScarlet puncta in the intestine (**Figures 1D-E**). Thus, hyperactivation of the p38 PMK-1 pathway by *vhp-1(RNAi)* phenocopied the TIR-1/SARM1 aggregation previously observed during pathogen infection.^12^

To test the hypothesis that p38 PMK-1 potentiates innate immune activation by increasing TIR-1/SARM1 expression, we crossed the p38/*pmk-1(km25)* null mutant into the strain expressing TIR-1::3xFLAG. The p38/*pmk-1(km25)* mutation suppressed induction of TIR-1::3xFLAG protein by *vhp-1(RNAi)* (**Figures 1A-B**). The phosphatase *vhp-1* also dephosphorylates the MAP kinase *kgb-1.*^15^ However, the *kgb-1(km21)* null allele did not suppress the induction of TIR-1::3xFLAG protein by *vhp-1(RNAi),* demonstrating that the p38 PMK-1 pathway specifically induced TIR-1/SARM1 in a feedforward cycle (**Figures S1B-C**).

Consistent with the TIR-1/SARM1 protein expression data, we also observed that knockdown of *vhp-1* induced the transcription of *tir-1* mRNA (**Figure 1F**). Importantly, the increased *tir-1* transcription observed in *vhp-1(RNAi)* animals was significantly suppressed in the p38/*pmk-1(km25*) mutant background (**Figure 1F**). Thus, activation of *tir-1* transcription by the p38 PMK-1 pathway potentiated innate immune signaling by inducing TIR-1/SARM1 multimerization. We propose that feedforward propagation of immune defenses in this manner facilitates a rapid and robust response to challenge by an infectious pathogen.

### A forward genetic screen identifies *rotr-1*, a novel suppressor of immune gene transcription

Constitutive activation of the p38 PMK-1 immune pathway is detrimental to the overall health of *C. elegans*.^16–20^ Thus, we reasoned that feedforward activation of p38 PMK-1 immune signaling is regulated to prevent the deleterious effects of unchecked innate immune activation. We performed a forward genetic screen to identify the mechanism that promotes this immune homeostasis. For these studies, we used a transcriptional reporter for the innate immune gene *irg-5* (*irg-5*p::*gfp*) to provide a visual readout of p38 PMK-1 pathway activation. The gene *irg-5* encodes a secreted innate immune effector whose transcription is strongly induced in intestinal epithelial cells during infection with several different bacterial pathogens, including *P. aeruginosa*.^10,12,21,22^ The basal expression of *irg-5* (*i.e.*, in the absence of pathogen infection) depends on the p38 PMK-1 pathway^10,23^ and knockdown of *irg-5* alone renders *C. elegans* hypersusceptible to pathogen infection.^23^ We therefore hypothesized that a genetic screen for mutants, which cause constitutive activation of *irg-5*, would uncover mechanisms that promote immune homeostasis in the *C. elegans* intestine.

Of 33,000 haploid mutant genomes screened from the F2 generation, nine mutant strains were recovered that hyperactivated *irg-5*p::*gfp* (**Figure S2A**). Three of these nine strains contained a mutation in Y42G9A.1 (*ums33, ums38, ums39*), a previously uncharacterized gene that we named regulator <u>o</u>f *<u>tir</u>-1* (*rotr-1)* (**Figure 2A**). Both *ums33* and *ums39* had a missense mutation converting serine to phenylalanine at position 92 (S92F), and *ums38* carried a nonsense mutation at position 186 (N186*) (**Figure 2A**). We used qRT-PCR to confirm that the native *irg-5* gene was hyperactivated in *rotr-1(ums33), rotr-1(ums38)* and *rotr-1(ums39)* loss-of-function mutants (**Figure 2B**). RNAi-mediated knockdown of *rotr-1* hyperactivated *irg-5*p::*gfp* (**Figure 2C**). Using CRISPR-Cas9, we generated a clean deletion mutant of *rotr-1* that spanned 4,621 bp of the *rotr-1* gene and 34 bp of the upstream promoter region [*rotr-1(ums53)*]. *C. elegans rotr-1(ums53)* mutants recapitulated the hyperactivation of *irg-5*p::*gfp* observed in the *rotr-1* mutants recovered in the forward genetic screen (**Figure 2D**). We raised an antibody to the ROTR-1 protein and confirmed that this protein was not expressed in *rotr-1(ums53)* mutants, demonstrating that this is a null allele (**Figure S2B**). Re-introduction of *rotr-1* expressed under the control of its own promoter into *rotr-1(ums38),* a mutant recovered in the forward genetic screen, suppressed the induction of *irg-5*p::*gfp* (**Figure 2E**).

**Figure 2.**
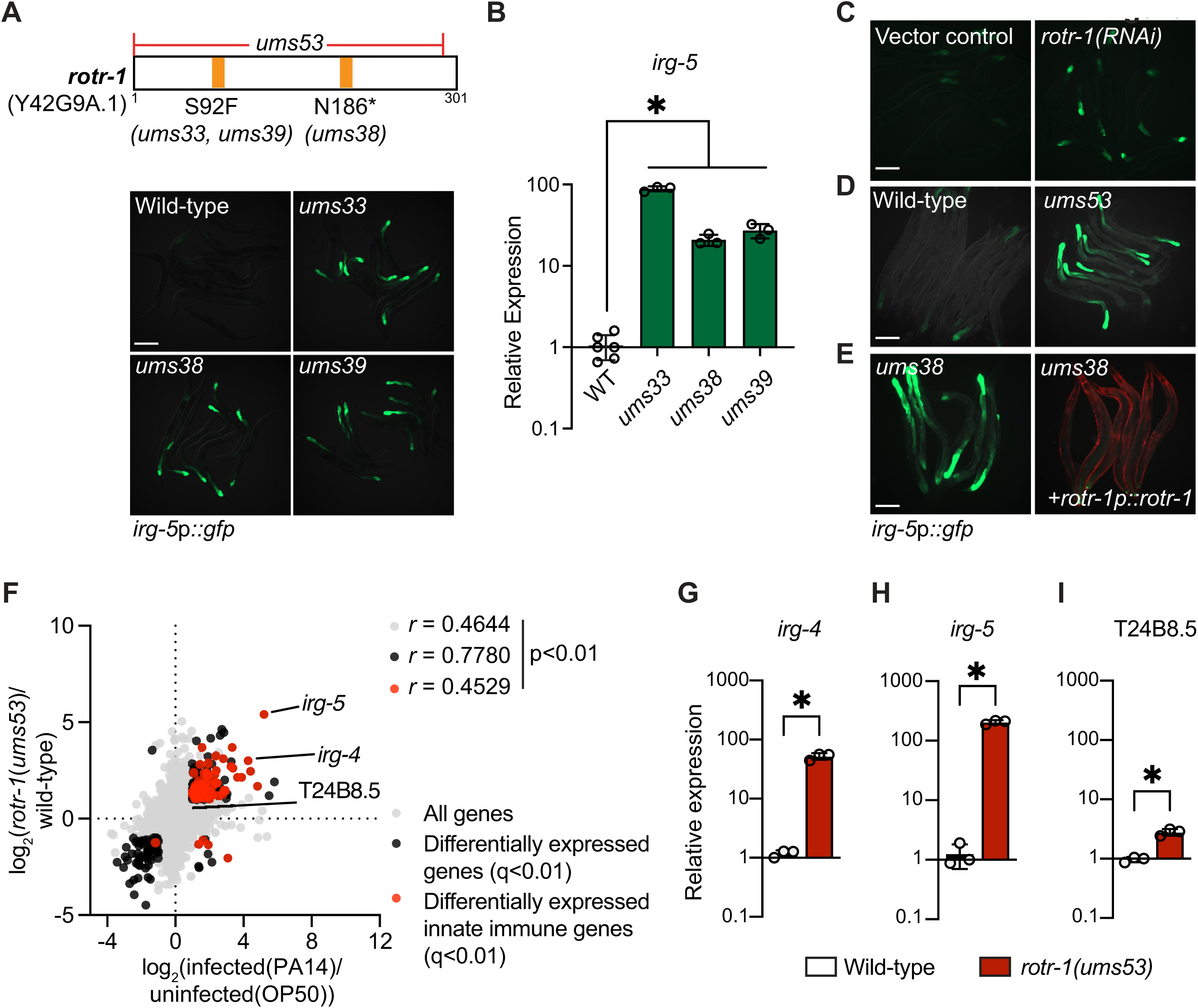
A forward genetic screen identifies *rotr-1*, a novel suppressor of immune gene transcription. **(A)** Representative images of forward genetic mutants *ums33, ums38,* and *ums39* in the *irg-5*p*::gfp* reporter background. Specific mutations of each allele are indicated in the schematic of the *rotr-1* protein sequence above the images. **(B)** qRT-PCR analysis of *irg-5* transcription in the forward genetic mutants in (A). *equals p<0.05 (one-way ANOVA with Dunnett’s multiple comparisons test). **(C)** Representative images of *irg-5*p::*gfp* animals treated with *rotr-1(RNAi).* **(D)** Representative images of wild-type and CRISPR-Cas9-generated *rotr-1(ums53)* clean deletion mutant in the *irg-5*p::*gfp* reporter background. **(E)** Representative images of *rotr-1(ums38),* a mutant isolated from the forward genetic screen that was rescued with a transgene expressing *rotr-1* under its own promoter, as indicated. Red indicates *myo-3*p*::mCherry* expression and is the co-injection marker. **(F)** Data from mRNA-seq experiments comparing genes differentially regulated in uninfected *rotr-1(ums53)* mutants versus wild-type animals or uninfected wild-type animals versus *P. aeruginosa-*infected wild-type animals. All genes are shown in gray. Genes that are differentially expressed in both datasets are shown in black (fold change>2, q<0.01). Genes that are annotated as innate immune genes are shown in red. The location of *irg-4, irg-5* and T24B8.5 is indicated. Note that T24B8.5 is only significantly differentiated in infected animals. See also Table S2. **(G-I)** qRT-PCR analyses comparing the transcription of *irg-4* (**G**), *irg-5* (**H**), and T24B8.5 (**I**) in wild-type and *rotr-1(ums53)* null mutants. *equals p<0.05 (unpaired t test). Source data for this figure is in Table S3. Scale bars represent 200 µm. See also Fig. S2.

mRNA-sequencing (seq) revealed that putative immune effectors were strongly upregulated in *rotr-1(ums53)* mutants (**Figure 2F**). We compared the genes that were differentially regulated in *rotr-1(ums53)* mutants with genes whose expression were changed in wild-type *C. elegans* during infection with *P. aeruginosa*. These datasets significantly correlated with each other (*r* = 0.778, p<0.01, black circles in **Figure 2F**). Intriguingly, the host defense genes that were differentially expressed in *rotr-1(ums53)* mutants were enriched in the upper right quadrant (red circles in **Figure 2F**), demonstrating that *rotr-1* suppressed the expression of putative immune effectors. Of note, *irg-5,* the transcriptional reporter used in the forward genetic screen, was the most significantly upregulated gene in *rotr-1(ums53)* mutants with a log_2_ fold-change of 5.4 compared to wild-type animals (**Figure 2F, Supplemental Table S2**). We confirmed the mRNA-seq data using qRT-PCR to assay the expression of three innate immune effectors in the *rotr-1(ums53)* clean deletion mutant: *irg-4* (**Figure 2G**); *irg-5* (**Figure 2H**); and T24B8.5 (**Figure 2I**).

In summary, a forward genetic screen identified *rotr-1*, a previously uncharacterized suppressor of immune gene transcription.

### ROTR-1 suppresses feedforward activation of p38 PMK-1 innate immunity

Gene set enrichment analysis of the RNA-seq data revealed that p38 PMK-1 targets were significantly enriched among the upregulated genes in *rotr-1(ums53)* (**Figure 3A**). Accordingly, *rotr-1(ums53)* activated a GFP transcriptional reporter for T24B8.5(*sysm-1*)(T24B8.5p::*gfp*), a putative ShK-like protein dependent on p38 PMK-1 (**Figure 3B**).^10,11^ To confirm that *rotr-1(ums53)* null mutants hyperactivated p38 PMK-1, we performed immunoblotting to quantify phosphorylated p38 PMK-1 compared to total p38 PMK-1. *C. elegans rotr-1(ums53)* null mutants had significantly higher levels of active p38 PMK-1 relative to wild-type animals (**Figure 3C-D**), consistent with the gene expression signature of these mutants. Importantly, the hyperphosphorylation of p38 PMK-1 in the *rotr-1(ums53)* mutants was suppressed in both the *tir-1(qd4)* and *pmk-1(km25)* loss-of-function backgrounds (**Figures 3C-D**). Consistent with these data, hyperactivation of the p38-dependent innate immune genes *irg-4* (**Figure S3A**)*, irg-5* (**Figure S3B**), and T24B8.5 (**Figure S3C**) in the *rotr-1(ums53)* null mutants was suppressed in the absence of *tir-1* or *pmk-1.* Thus, ROTR-1 suppressed p38 PMK-1 pathway activation.

**Figure 3.**
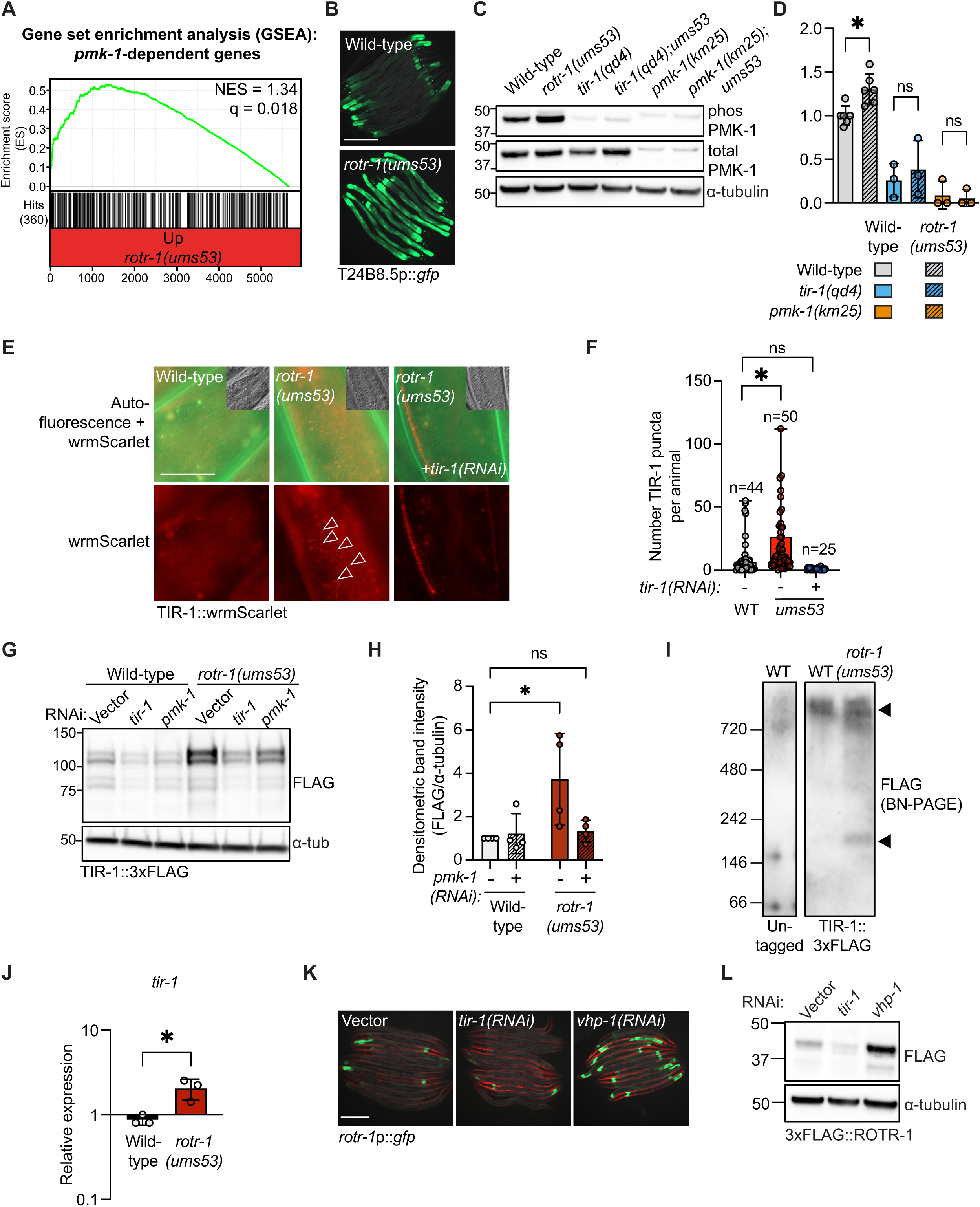
ROTR-1 suppresses feedforward activation of p38 PMK-1 innate immunity. **(A)** Gene set enrichment analysis (GSEA) of p38 PMK-1 targets in the *rotr-1(ums53)* mRNA-seq experiment. Each differentially upregulated gene in *rotr-1(ums53)* mutants was assigned a π-value^49^ and ranked from highest to lowest. Normalized enrichment score (NES) and q-value are indicated. p38 PMK-1 targets identified are indicated by hit number and black lines below the graph. **(B)** Representative images of wild-type and *rotr-1(ums53)* null mutants in the T24B8.5p::*gfp* transcriptional reporter animals. **(C)** Immunoblot of whole cell lysates from indicated genotypes that were probed for anti-phosphorylated PMK-1, anti-total PMK-1, and anti-α-tubulin. **(D)** Densitometric quantification of (C). Error bars represent SEM. *equals p<0.05 (two-way ANOVA with Šídák’s multiple comparisons test). **(E)** Images of wild-type or *rotr-1(ums53)* mutants expressing TIR-1::wrmScarlet. Insets represent corresponding DIC images. Arrowheads indicate TIR-1::wrmScarlet puncta. **(F)** Quantification of TIR-1::wrmScarlet puncta in (E) as previously described.^12^ *equals p<0.05 (one-way ANOVA with Dunnett’s multiple comparisons test). ns=not significant. **(G)** Immunoblot of whole cell lysates from wild-type and *rotr-1(ums53)* in the TIR-1::3xFLAG background treated with indicated RNAi and probed with anti-FLAG or anti-α-tubulin. **(H)** Densitometric quantification of vector control and *pmk-1(RNAi)* conditions in (G). *equals p<0.05 (two-way ANOVA with Šídák’s multiple comparisons test). **(I)** Blue NativePage immunoblot on wild-type and *rotr-1(ums53)* mutants in the TIR-1::3xFLAG background. Multimers are indicated by the arrowheads. Immunoblot of untagged animals is shown on the left panel. Immunoblot was probed using the anti-FLAG antibody. **(J)** qRT-PCR analysis of *tir-1* transcription in wild-type and *rotr-1(ums53)* mutants. *equals p<0.05 (unpaired t-test). **(K)** Representative images of animals expressing a *gfp* transcriptional fusion under the control of the *rotr-1* promoter (*rotr-1*p*::gfp)* treated with vector control, *tir-1(RNAi),* or *vhp-1(RNAi)*. (**L)** Immunoblot of whole cell lysates from 3xFLAG::ROTR-1 animals treated with vector control, *tir-1(RNAi)* or *vhp-1(RNAi).* Scale bars equal 200 µm. Source data for this figure is in Table S3. See also Fig. S3.

*C. elegans rotr-1(ums53)* mutants had significantly more TIR-1::wrmScarlet puncta than wild-type animals (**Figure 3E-F**). To determine whether TIR-1 protein levels were also increased in *rotr-1* mutants, we assessed TIR-1::3xFLAG protein expression by immunoblot. TIR-1::3xFLAG protein levels were significantly higher in *rotr-1(ums53)* mutants compared to wild-type animals (**Figure 3G-H**). In addition, *rotr-1(ums53)* null mutants expressed more higher-order multimers of TIR-1/SARM1 compared to wild-type animals when examined under native conditions (**Figure 3I**). Thus, our data demonstrated that ROTR-1 acts upstream of TIR-1/SARM1 to suppress p38 PMK-1 pathway activation.

Importantly, the increase in TIR-1/SARM1 protein in *rotr-1(ums53)* mutants was suppressed by RNAi-mediated knockdown of p38 *pmk-1* (**Figure 3G-H**), which recapitulated our previous observations with *vhp-1(RNAi)* (**Figure 1**). In addition, *rotr-1(ums53)* mutants expressed significantly higher levels of *tir-1* mRNA compared to wild-type animals (**Figure 3J**). Thus, ROTR-1 suppressed the feedforward activation of p38 PMK-1.

We generated a *gfp*-based transcriptional immune reporter for *rotr-1* to study its regulation. Knockdown of *vhp-1*, the phosphatase that negatively regulates p38 PMK-1, activated *rotr-1*p::*gfp* transcription (**Figure 3K**). In contrast, *tir-1*(RNAi) suppressed *rotr-1*p::*gfp* transcription (**Figure 3K**). Using CRISPR-Cas9, we introduced a 3xFLAG tag at the endogenous N-terminus of ROTR-1. As we observed in studies of *rotr-*1p::*gfp* transcription, *vhp-1(RNAi)* increased and *tir-1(RNAi)* suppressed 3xFLAG::ROTR-1 protein expression (**Figure 3L**). Thus, p38 PMK-1 pathway activation increased the expression of ROTR-1, a negative regulator of immune pathway signaling.

High-throughput interactome mapping of *C. elegans* proteins using yeast two-hybrid experiments previously found that ROTR-1 binds to TIR-1/SARM1 *in vitro*.^24^ These data raise the intriguing hypothesis that ROTR-1 physically restrains TIR-1/SARM1 to suppress p38 PMK-1 activation. We therefore sought to determine if this protein-protein interaction occurs *in vivo.* Using CRISPR-Cas9, we introduced a 3xHA tag at the C-terminus of ROTR-1 in an animal expressing TIR-1::3xFLAG. Unexpectedly, addition of the endogenous ROTR-1::3xHA tag induced TIR-1::3xFLAG protein expression (**Figure S3D**), phenocopying the upregulation of TIR-1/SARM1 protein expression in *rotr-1(ums53)* null mutants (**Figures 3G-H**). To test whether this result was terminus-specific, we used CRISPR-Cas9 to create two different strains that carried either a 3xFLAG or a 3xHA tag at the N-terminus of ROTR-1. However, the addition of these tags to ROTR-1 caused hyperactivation of the T24B8.5p::*gfp* reporter, which also phenocopied the *rotr-1(ums53)* mutation (**Figures S3E-F**). We conclude that tagging ROTR-1 at either the N- or C-termini disrupted the function of this protein, which precluded the ability to interpret data from co-immunoprecipitation experiments. Our attempts to use the antibody we raised against ROTR (**Figure S2B**) in co-immunoprecipitation experiments with TIR-1::3xFLAG were also unsuccessful secondary to non-specific protein binding by the ROTR-1 antibody. Nevertheless, these data provided additional confirmation of the phenotypes in the *rotr-1* loss-of-function mutant strain.

In summary, ROTR-1 suppressed TIR-1/SARM1 aggregation to restrain the feedforward propagation of p38 PMK-1 immune signaling.

### ROTR-1 functions in the intestine to suppress p38 PMK-1 innate immunity and support *C. elegans* growth and longevity

The *gfp*-based transcriptional reporter we generated for *rotr-1* revealed that this gene is expressed exclusively in the intestine, particularly in anterior and posterior intestinal epithelial cells (**Figure 3K**). To determine if *rotr-1* also functions in the intestine to regulate the p38 PMK-1 pathway, we engineered *C. elegans* strains that expressed *rotr-1* under the control of promoters that directed its expression only in specific tissues. Endogenous expression of *rotr-1* under its own promoter suppressed the hyperactivation of T24B8.5p::*gfp* in the *rotr-1(ums53)* clean deletion mutant (**Figure 4A**), as we observed for *irg-5*p::*gfp* in the rescued *rotr-1(ums38)* mutant (**Figure 2E**). Intestinal expression of *rotr-1* (**Figure 4B**), but not neuronal (**Figure 4C**) or hypodermal expression (**Figure 4D**), suppressed the hyperactivation of T24B8.5p::*gfp* expression in *rotr-1(ums53)* mutants. We confirmed the tissue-specific expression of *rotr-1* in these strains using a construct that contains a split-leader mCherry sequence, which labeled the tissues with *rotr-1* expression (**Figures 4B-D**).

**Figure 4.**
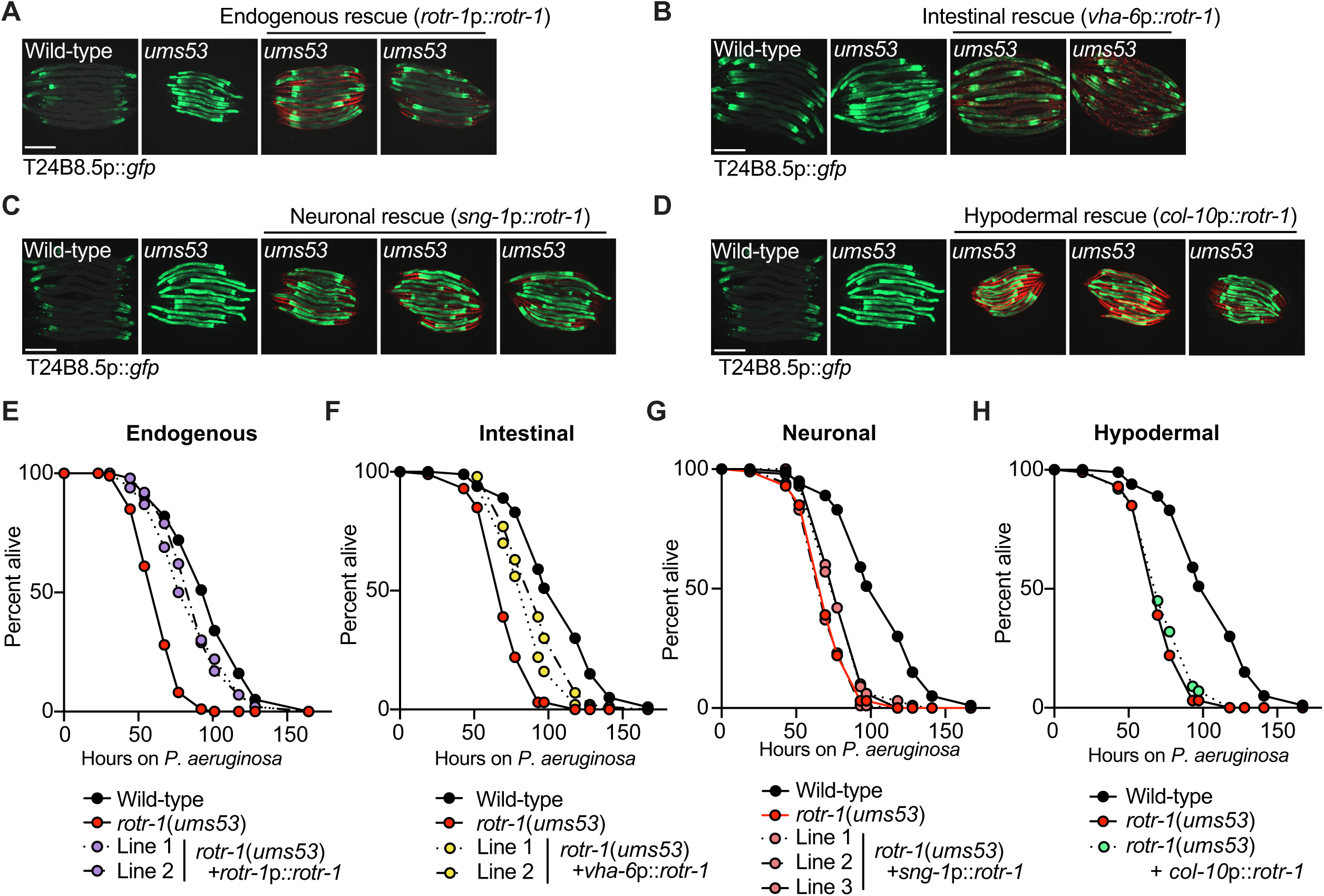
ROTR-1 functions in the intestine of *C. elegans*. **(A-D)** Images of wild-type and *rotr-1(ums53)* mutants expressing transgenes of *rotr-1* under the indicated promoters for endogenous *rotr-1* expression (*rotr-1*p*::rotr-1)* **(A)**, intestinal *rotr-1* expression (*vha-6*p::*rotr-1*) **(B)**, neuronal *rotr-1* expression (*sng-1*p::*rotr-1*) **(C)**, and hypodermal expression (*col-10*p::*rotr-1*) **(D)**. Tissue-specific expression of *rotr-1* in these strains was confirmed using a construct that contains a split-leader mCherry sequence for *his-58*, which labeled the nuclei of tissues with *rotr-1* expression in red. Source data for this figure is in Table S3. **(E-H)** Representative *P. aeruginosa* pathogenesis assays for strains indicated in (A-D). Note that only one hypodermal *rotr-1* rescue line is represented due to the toxicity of hypodermal expression of *rotr-1.* The difference between each rescue line and the *rotr-1(ums53)* mutant is significant in (E-F) (p<0.05, log-rank test). Scale bars equal 200 µm. Sample size (n), mean lifespans and statistics for all replicates are in Table S1.

We found that *rotr-1* loss-of-function mutants were hypersusceptible to killing by *P. aeruginosa* infection (**Figure 4E**), as we have observed with other mutants that also hyperactivate the p38 PMK-1 pathway.^12,16^ Endogenous rescue of *rotr-1* under its own promoter restored resistance of *rotr-1(ums53)* null mutants against killing by *P. aeruginosa* (**Figure 4E**). Reintroduction of *rotr-1* in the intestine (**Figure 4F**), but not in the neurons (**Figure 4G**) or hypodermis (**Figure 4H**), restored wild-type resistance to pathogen infection.

Our laboratory and others have previously shown that aberrant activation of the p38 PMK-1 pathway is deleterious to *C. elegans* growth and lifespan.^12,16,17,20,25,26^ Consistent with these studies, *rotr-1(ums53)* mutants had a significantly reduced lifespan (**Figure 5A**) and delayed development (**Figure 5B-C**). Importantly, each of these defects in *rotr-1(ums53)* mutants was rescued in a *tir-1(qd4)* loss-of-function background (**Figure 5A-C**), demonstrating that these effects were due to toxicities caused by hyperactivation of TIR-1/SARM1.

**Figure 5.**
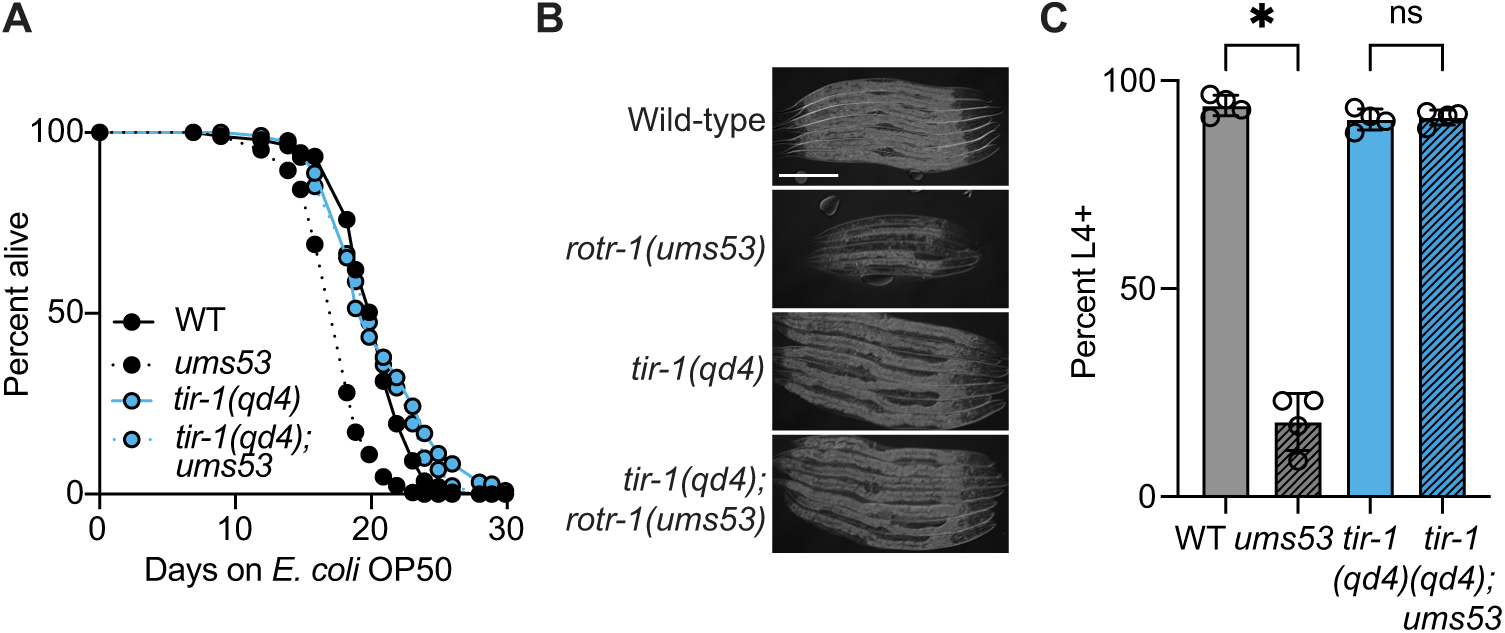
ROTR-1 is required for *C. elegans* growth and longevity. **(A)** Representative lifespan assay of wild-type, *rotr-1(ums53), tir-1(qd4)* and *tir-1(qd4);rotr-1(ums53)* mutants. **(B)** DIC images of indicated *C. elegans* genotypes in a development assay. Scale bar equals 200 µm. **(C)** Development assay of indicated *C. elegans* genotypes quantifying the percent of animals older than the larval L4 stage. *equals p<0.05, ns=not significant (one-way ANOVA with Dunnett’s multiple comparisons test). Sample size (n), mean lifespans and statistics for all replicates are in Table S1.

In summary, *rotr-1* functions in the intestine to restrain toxic activation of TIR-1/SARM1.

### ROTR-1 supports the integrity of lysosome-related organelles, which restrains feedforward activation of p38 PMK-1 innate immunity

In a companion manuscript, we discovered that the conserved signaling regulator TIR-1/SARM1, the upstream activator of the p38 PMK-1 signaling cassette, is expressed on the membranes of a specific population of lysosomes called lysosome-related organelles. Aggregation of TIR-1/SARM1 into puncta following the pathogen-induced condensation of lysosome-related organelles engages the intrinsic NAD^+^ hydrolase activity of this protein complex to activate p38 PMK-1 innate immune defenses.

In the mRNA-seq experiment (**Figure 2**), we also found that *rotr-1* regulated a significant number of genes involved in lysosomal function, including acid phosphatases and genes required for proteolysis (**Figure S4, Table S2**). In light of the findings presented in our companion manuscript, we hypothesized that ROTR-1 promotes the integrity of lysosomes, which in turn restrains feedforward activation of p38 PMK-1 innate immunity. To test whether *rotr-1(ums53)* null mutants had defects in the lysosomal compartment, we used LysoTracker Red, a dye that stains acidic organelles.^27,28^ Compared to wild-type animals, *rotr-1(ums53)* mutants had significantly fewer vesicles that stained positively for LysoTracker Red, demonstrating that the lysosomal compartment is compromised in this mutant background (**Figure 6A-B**). Importantly, in our companion manuscript, we discovered that both *P. aeruginosa* infection and treatment with the secreted pseudomonal virulence effector pyocyanin depleted Lysotracker Red staining of acidic vesicles in intestinal epithelial cells. Thus, the *rotr-1(ums53)* mutant recapitulates the changes in the lysosomal compartment that are observed during pathogen infection.

**Figure 6.**
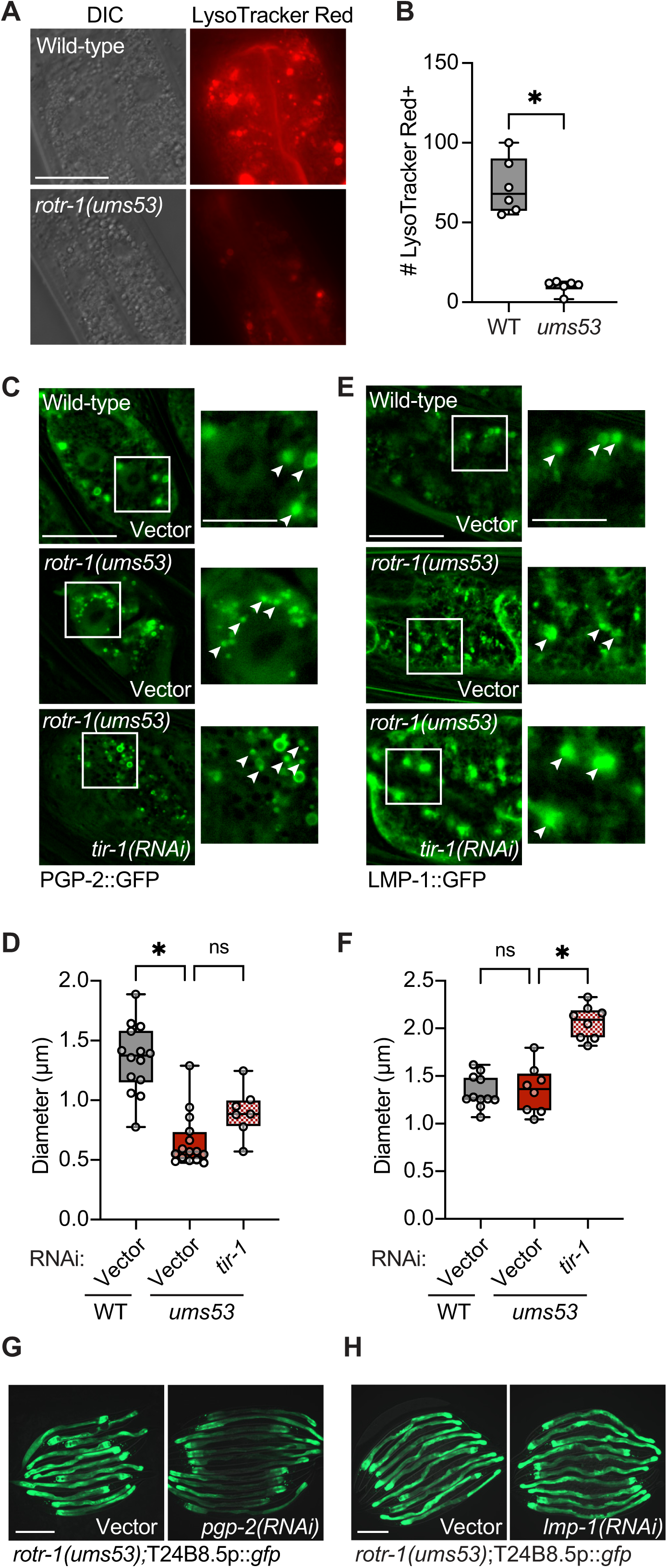
ROTR-1 supports the integrity of lysosome-related organelles, which restrains feedforward activation of p38 PMK-1 innate immunity. **(A)** Images of wild-type and *rotr-1(ums53)* mutants stained with LysoTracker Red (1 µM). DIC images are shown on the left. Scale bar equals 20 µm. **(B)** Quantification of the number of LysoTracker Red(+) vesicles in wild-type and *rotr-1(ums53)* mutants using Fiji (*n=*6). **(C)** Images of wild-type and *rotr-1(ums53)* mutants expressing PGP-2::GFP (lysosome-related organelles) treated with vector control or *tir-1(RNAi)*. Boxes indicate area shown at higher magnification to the right. Scale bar equals 20 µm for lower magnification and 10 µm for higher magnification. **(D)** Quantification of the diameter of PGP-2::GFP(+) vesicles in indicated conditions. Each data point represents one animal. *equals p<0.05, ns=not significant (one-way ANOVA with Tukey’s multiple comparisons test). **(E)** Images of wild-type and *rotr-1(ums53)* mutants expressing LMP-1::GFP (lysosomes) treated with vector control or *tir-1(RNAi)*. Boxes indicate area shown at higher magnification to the right. Scale bar equals 20 µm for lower magnification and 10 µm for higher magnification. **(F)** Quantification of the diameter of LMP-1::GFP(+) vesicles in indicated conditions. *equals p<0.05, ns=not significant (one-way ANOVA with Tukey’s multiple comparisons test). **(G-H)** Representative images of *rotr-1(ums53)* mutants in the T24B8.5p::*gfp* background treated with vector control or *pgp-2(RNAi)* (G) or *lmp-1(RNAi)* (H). Scale bar equals 200 µm. See Table S3. See also Figure S4.

Both lysosomes and lysosome-related organelles stain positively for LysoTracker Red in *C. elegans* intestinal tissues.^27–29^ To determine which of these cellular compartments is affected in *rotr-1(ums53)* mutants, we used transgenic *C. elegans* strains with GFP markers specifically labeled for either lysosomes or lysosome-related organelles. For lysosomes, we used a GFP translational fusion for the protein LMP-1 (LMP-1::GFP), which is the *C. elegans* homolog of mammalian LAMP and CD68.^30^ To label lysosome-related organelles, we used a GFP translational fusion for the protein PGP-2 (PGP-2::GFP), an ATP-binding cassette transporter that is expressed on the membranes of these vesicles.^27^ Intriguingly, the size of PGP-2::GFP (+) vesicles (**Figures 6C-D**), but not LMP-1::GFP (+) vesicles (**Figures 6E-F**), were significantly smaller in *rotr-1(ums53)* mutants compared to wild-type animals. Thus, the integrity of lysosome-related organelles, but not lysosomes, was compromised in *rotr-1(ums53)* mutants. This result is noteworthy in light of our findings that TIR-1/SARM1 is expressed on the membranes of lysosome-related organelles, but not on lysosomes (companion manuscript).

Importantly, knockdown of *tir-1* did not rescue the collapsed PGP-2::GFP (+) vesicles in the *rotr-1(ums53)* mutants (**Figure 6C-D**). Thus, the condensation of lysosome-related organelles in *rotr-1* mutants occurred upstream of TIR-1/SARM1. These data are important considering the findings in our companion study, which demonstrated that TIR-1/SARM1 aggregates into puncta on the surface of condensed lysosome-related organelles to activate the p38 PMK-1 pathway. Consistent with these observations, the hyperactivation of the p38 PMK-1-dependent immune reporter in *rotr-1(ums53)* mutants was dependent on the presence of lysosome-related organelles. RNAi-mediated knockdown of *pgp-2* (**Figure 6G**), but not *lmp-1* (**Figure 6H**) suppressed T24B8.5p::*gfp* activation in the *rotr-1(ums53)* mutants.

Together, these data demonstrated that lysosomal integrity, which was compromised in *rotr-1* mutants, restrained toxic feedforward activation of p38 PMK-1 innate immunity by preventing aberrant TIR-1/SARM1 aggregation.

## DISCUSSION

Enzymatic TIR proteins have essential roles in innate immunity in bacteria, plants, and animals. Across the tree of life, members of this protein family oligomerize to initiate their intrinsic NAD^+^ catalytic activity.^3,5,7,31,32^ However, the mechanisms that control TIR protein multimerization are under-explored. In this study, we showed that the activity of TIR-1/SARM-1, the lone enzymatic TIR protein in *C. elegans*, is controlled through the maintenance of lysosome-related organelle integrity. Initially, we discovered that TIR-1/SARM1 promotes its own activation by initiating a p38 PMK-1-dependent positive feedforward loop, which drives the transcription, translation, and aggregation of TIR-1/SARM1. Unchecked TIR-1/SARM1 aggregation and activation decreased lifespan, reduced brood size, and stunted larval development. The previously uncharacterized protein ROTR-1 functions in intestinal tissues to maintain the integrity of the specific lysosomal compartment that expresses TIR-1/SARM1, which restrains p38 PMK-1 innate immune activation by preventing aberrant TIR-1/SARM1 aggregation.

In a companion study, we showed that TIR-1/SARM1 is a pathogen sensor in animals, functioning to initiate p38 innate immune activation from the surface of the specific lysosomal compartment whose integrity is ensured by ROTR-1. A redox active virulence effector secreted by the bacterial pathogen *Pseudomonas aeruginosa* alkalinized and collapsed lysosome-related organelles, which promoted TIR-1/SARM1 aggregation on their surface. Together, our companion studies demonstrate that the lysosomal – TIR-1/SARM1 – p38 axis functions in the pathogen effector-triggered activation of protective host immunity and is tightly regulated to promote intestinal immune homeostasis.

We propose that p38 feedforward potentiation of immune defenses promotes a rapid host response to pathogen infection. Concentration of TIR-1/SARM1 into puncta, or higher-order oligomers, accelerates its enzymatic function.^8,12^ It is therefore logical that feedforward immune activation in *C. elegans* is caused by increasing the transcription and translation of TIR-1/SARM1, which increases the likelihood of protein oligomerization. Perhaps not surprisingly, we found that excessive TIR-1/SARM1 activity is detrimental to overall animal health. These data are consistent with findings from our group and others, which have demonstrated that chronic hyperactivation of the p38 PMK-1 pathway is toxic to nematodes.^16,33,34^ Here, we demonstrated that these immune toxicities are controlled at the level of TIR-1/SARM1 oligomerization.

SARM1 is expressed in neurons where it promotes Wallerian degeneration of axon fragments following neuronal injury.^8,35–39^ In this regard, SARM1 has emerged as an attractive drug target to prevent pathologic neurodegeneration.^40^ *C. elegans* TIR-1/SARM1 has also been implicated in neuronal degeneration.^8,41^ In this context, overexpression of TIR-1/SARM1 promotes toxic axonal degeneration; however, what causes this toxicity is not known. Thus, an intriguing hypothesis is that lysosomal regulation of TIR-1/SARM1 aggregation in neurons may mitigate TIR-1/SARM1-induced neurotoxicity.

We demonstrated that ROTR-1 is required for the integrity of lysosome-related organelles, but it is not clear how ROTR-1 functions in this capacity. Our efforts to characterize the cell biology of ROTR-1 were unsuccessful given the inability to generate a functional tagged protein. Thus, we do not know if ROTR-1 is expressed on lysosome-related organelles, nor could we confirm *in vivo* the previous observation that ROTR-1 physically interacts with TIR-1/SARM1. Nevertheless, our studies with *rotr-1* mutants characterized the importance of this specific cellular compartment in the regulation of the p38 PMK-1 immune pathway.

It is important to note that p38 PMK-1 promotes host adaptation to a multitude of exogenous and endogenous stresses.^10,12,14,17,18,21,42–47^ In this regard, multiple different mechanisms have been identified that control the activity of this critically important cytoprotective kinase. The MAPK phosphatase *vhp-1* is a negative regulator of p38 PMK-1.^14^ Several different amphid sensory neurons send signals to the intestine to suppress p38 PMK-1 activity.^17,48^ Micronutrient deficiency primes p38 PMK-1 activation as part of an adaptive response to anticipate environmental threats during a time of relative vulnerability.^12^ We propose here that each of these different mechanisms of p38 PMK-1 regulation are linked to cell biological mechanisms that are perturbed by the specific stressors that are surveilled to activate protective host defenses. In the case of the bacterial effector-triggered p38 immune activation induced by lysosomal damage, we demonstrated that the lysosomal compartment that expresses the activator of p38 is itself regulated to restrain exaggerated immune responses. Thus, we propose that mechanisms of pathogen detection – particularly effector-triggered immunity – evolved specific protective countermeasures to prevent unchecked immune activation.

## ACKNOWLEDGEMENTS

The authors thank Melanie Trombly for insightful comments on the manuscript. The research was supported by R01 AI130289 (to R.P.W.), R01 AI159159 (to R.P.W.), R21 AI163430 (to R.P.W.), F30 DK127690 (to S.Y.T.), T32 AI095213 (to S.Y.T.), and T32 GM107000 (to S.Y.T.). Some strains were provided by the *Caenorhabditis* Genetics Center, which is funded by the NIH Office of Research Infrastructure Programs (P40 OD010440). The funders had no role in study design, data collection and analysis, decision to publish, or preparation of the manuscript.

## AUTHOR CONTRIBUTIONS

Conceptualization: SYT, RPW; Methodology: SYT, RPW; Investigation: SYT; Visualization: SYT; Funding acquisition: RPW; Project administration: RPW; Supervision: RPW; Writing – original draft, SYT, RPW; Writing – review & editing, SYT, RPW

## INCLUSION AND DIVERSITY STATEMENT

We support inclusive, diverse, and equitable conduct of research.

## STAR METHODS

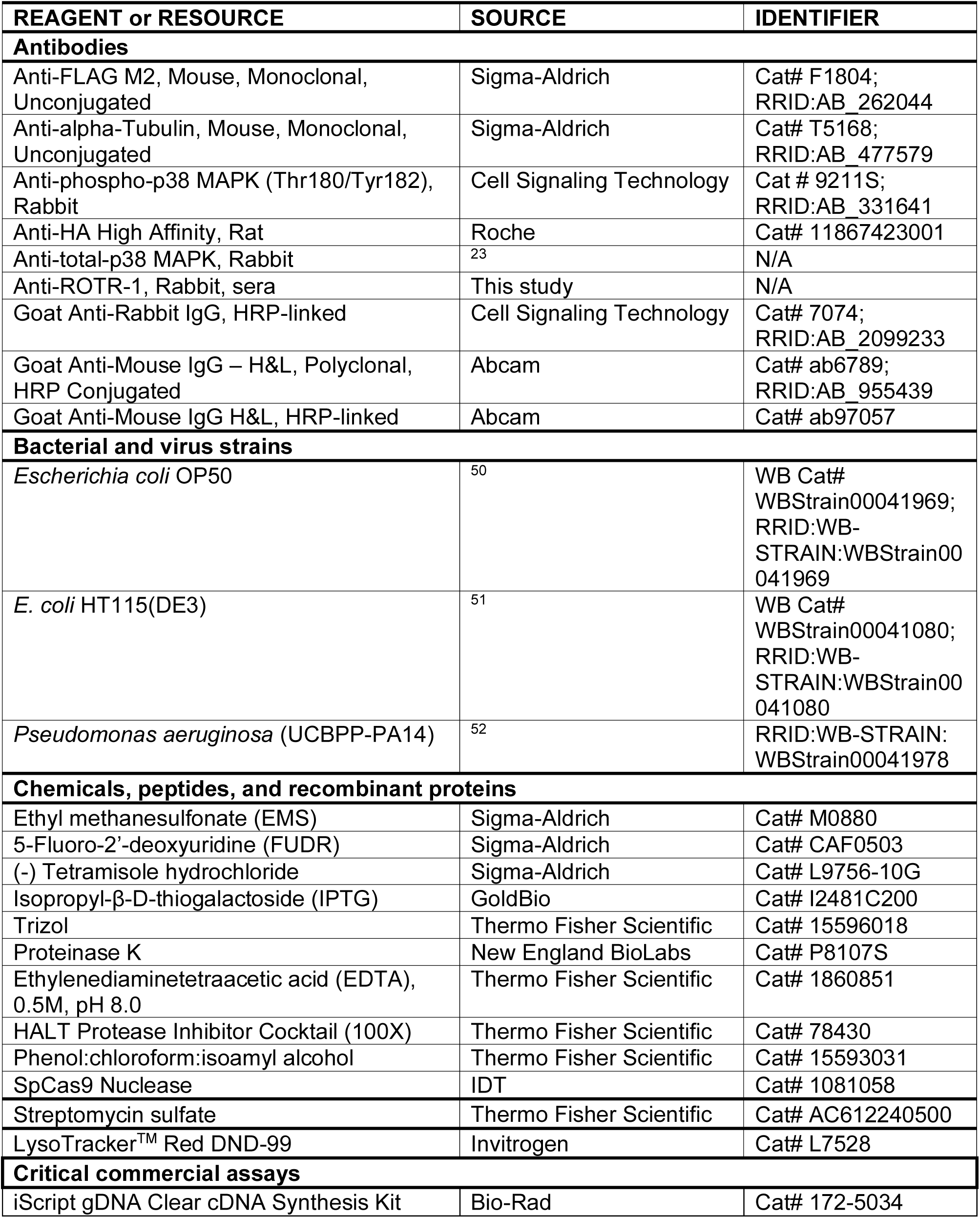
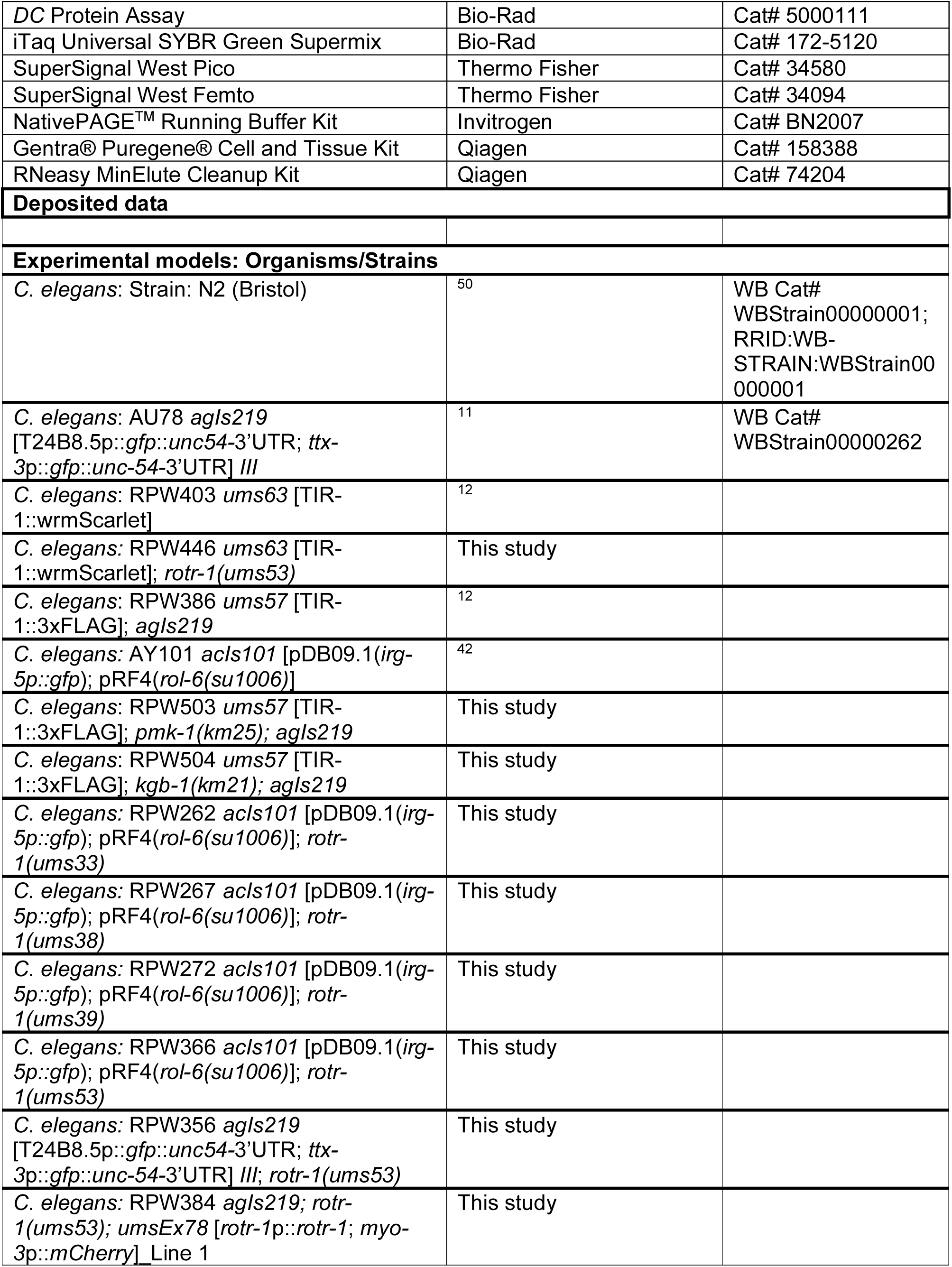
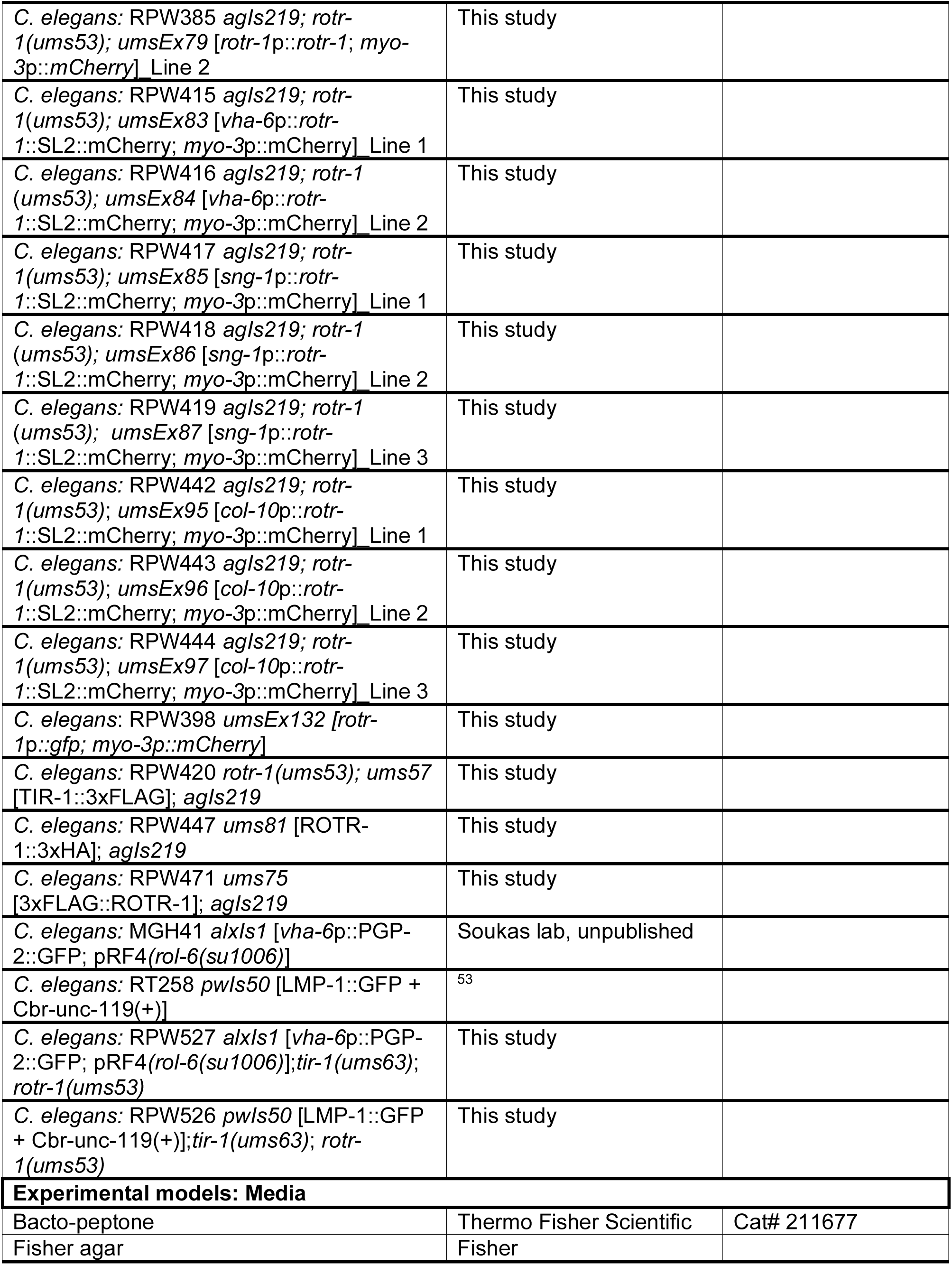
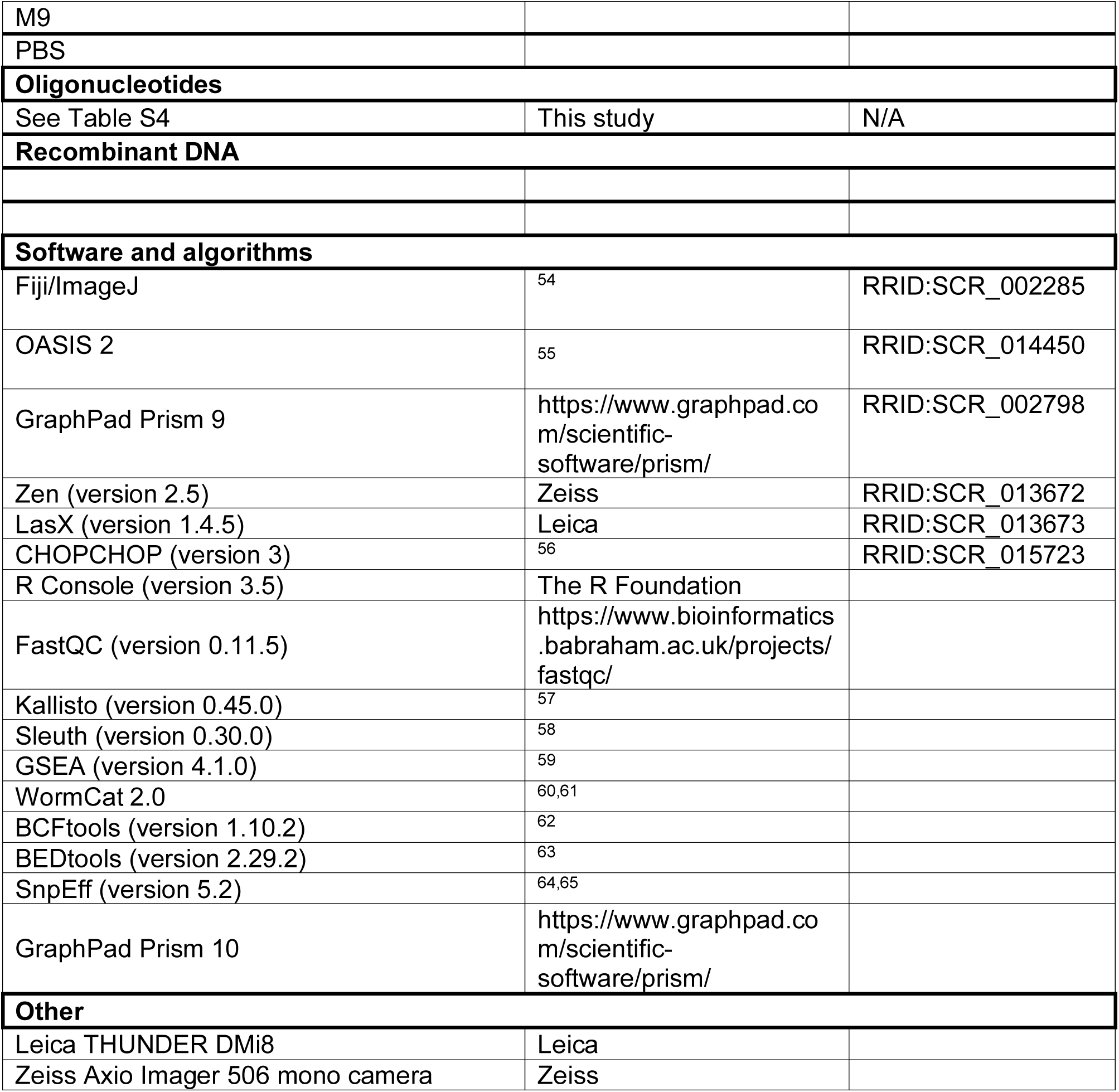
KEY RESOURCES TABLE.

## RESOURCE AVAILABILITY

### Lead contact

Further information and requests for resources or reagents should be directed to and will be fulfilled by the lead contact, Read Pukkila-Worley.

### Material availability

Strains and reagents generated in this study are available upon request.

### Data and code availability

- The mRNA-seq datasets will be deposited at NCBI Gene Expression Omnibus and will be publicly available as of the date of publication. All other data are available in the manuscript and the accompanying Table S3, which contains all source data and statistical tests used.
- This paper does not report original code.
- Any additional information required to reanalyze the data reported in this paper is available from the lead contact upon request.

## EXPERIMENTAL MODEL AND SUBJECT DETAILS

### C. elegans strains

The previously published *C. elegans* strains used in this study were: N2 Bristol,^50^ AU78 *agIs219* [T24B8.5p::*gfp*::*unc54-*3’UTR; *ttx-3*p::*gfp*::*unc-54-*3’UTR] *III,*^11^ RPW403 *ums63* [TIR-1::wrmScarlet],^12^ RPW386 *ums57* [TIR-1::3xFLAG]; *agIs219* [T24B8.5p::*gfp*::*unc54-*3’UTR; *ttx-3*p::*gfp*::*unc-54-*3’UTR] *III,*^12^ MGH41 *alxIs1* [*vha-6*p::PGP-2::GFP;pRF4] (unpublished), RT258 *pwIs50* [LMP-1::GFP + Cbr-unc-119(+)].^53^

The strains developed in this study were: RPW446 *ums63* [TIR-1::wrmScarlet]; *rotr-1(ums53),* RPW262 *acIs101* [pDB09.1(*irg-5p::gfp*); pRF4(*rol-6(su1006)*]; *rotr-1(ums33),* RPW267 *acIs101* [pDB09.1(*irg-5p::gfp*); pRF4(*rol-6(su1006)*]; *rotr-1(ums38),* RPW272 *acIs101* [pDB09.1(*irg-5p::gfp*); pRF4(*rol-6(su1006)*]; *rotr-1(ums39),* RPW366 *acIs101* [pDB09.1(*irg-5p::gfp*); pRF4(*rol-6(su1006)*]; *rotr-1(ums53),* RPW356 *agIs219* [T24B8.5p::*gfp*::*unc54-*3’UTR; *ttx-3*p::*gfp*::*unc-54-* 3’UTR] *III*; *rotr-1(ums53),* RPW384 *agIs219; rotr-1(ums53); umsEx78* [*rotr-1*p::*rotr-1*; *myo-3*p::*mCherry*]_Line 1, RPW385 *agIs219; rotr-1(ums53); umsEx79* [*rotr-1*p::*rotr-1*; *myo-3*p::*mCherry*]_Line 2, *C. elegans:* RPW415 *agIs219; rotr-1*(*ums53); umsEx83* [*vha-6*p::*rotr-1*::SL2::mCherry; *myo-3*p::mCherry]_Line 1, RPW416 *agIs219; rotr-1* (*ums53); umsEx84* [*vha-6*p::*rotr-1*::SL2::mCherry; *myo-3*p::mCherry]_Line 2, RPW417 *agIs219; rotr-1(ums53); umsEx85* [*sng-1*p::*rotr-1*::SL2::mCherry; *myo-3*p::mCherry]_Line 1, RPW418 *agIs219; rotr-1* (*ums53); umsEx86* [*sng-1*p::*rotr-1*::SL2::mCherry; *myo-3*p::mCherry]_Line 2, RPW419 *agIs219; rotr-1* (*ums53); umsEx87* [*sng-1*p::*rotr-1*::SL2::mCherry; *myo-3*p::mCherry]_Line 3, RPW442 *agIs219; rotr-1(ums53)*; *umsEx95* [*col-10*p::*rotr-1*::SL2::mCherry; *myo-3*p::mCherry]_Line 1, RPW443 *agIs219; rotr-1(ums53)*; *umsEx96* [*col-10*p::*rotr-1*::SL2::mCherry; *myo-3*p::mCherry]_Line 2, RPW444 *agIs219; rotr-1(ums53)*; *umsEx97* [*col-10*p::*rotr-1*::SL2::mCherry; *myo-3*p::mCherry]_Line 3, RPW398 *umsEx132 [rotr-1*p*::gfp; myo-3p::mCherry*], RPW420 *rotr-1(ums53); ums57* [TIR-1::3xFLAG]; *agIs219,* RPW447 *ums81* [ROTR-1::3xHA]; *agIs219,* RPW471 *ums75* [3xFLAG::ROTR-1]; *agIs219,* RPW527 *alxIs1* [*vha-6*p::PGP-2::GFP; pRF4*(rol-6(su1006)*];*tir-1(ums63)*; *rotr-1(ums53),* and RPW526 *pwIs50* [LMP-1::GFP + Cbr-unc-119(+)];*tir-1(ums63)*; *rotr-1(ums53)*.

### *C. elegans* growth conditions

*C. elegans* strains were maintained on standard nematode growth medium (NGM) plates [0.25% Bacto-peptone, 0.3% sodium chloride, 1.7% agar (Fisher agar), 5 μg/mL cholesterol, 25 mM potassium phosphate pH 6.0, 1 mM magnesium sulfate, 1 mM calcium chloride] with *E. coli* OP50 as a food source, as described.^50^

### Bacterial strains

Bacteria used in this study were *Escherichia coli* (*E. coli*) OP50,^50^ *E. coli* HT115(DE3),^51^ and *Pseudomonas aeruginosa* strain PA14.^52^

### Bacterial growth conditions

*E. coli* OP50 were grown in LB broth supplemented with 0.175 mg/mL streptomycin at 37 °C for 16-18 hrs at 250 rpm. *P. aeruginosa* strains were grown in LB broth at 37 °C for 14-15 hrs at 250 rpm.

## METHODS

### Identification of *rotr-1* through an unbiased *irg-5* forward genetic screen

Ethyl methanesulfonate (EMS) mutagenesis was performed on strain *acIs101* (*irg-5*p*::gfp)* as previously described.^16,17^ Briefly, synchronized L4 animals were treated with 48.6 mM EMS in M9 liquid for 4 hrs at 22°C on a roller. P0 animals were plated onto NGM plates with *E. coli* OP50. Gravid F1 progeny were then treated with hypochlorite and eggs were allowed to hatch overnight. Synchronized F2 progeny were grouped into 7 genetically distinct pools and screened for bright and constitutive GFP expression. Approximately 33,000 haploid genomes from the F2 generation were screened. Nine mutants were recovered. Animals with significant developmental delay were excluded. From this screen, 9 mutants were identified. Forward genetic mutants *ums33* and *ums39* came from the same pool and *ums38* came from a separate pool.

Mutants were backcrossed with the parent strain (*acIs101*) two times. Progeny from each mutant, unbackcrossed and backcrossed, were then pooled. Genomic DNA from the pooled recombinants and parent strain were isolated using the Gentra® Puregene® DNA isolation kit (Qiagen) and sent for whole genome sequencing (BGI). Animals were sequenced on the DNBseq platform with 100 bp paired-end runs. Each sample had an average of 65 million reads with a final coverage of around 130x.

Homozygous variants from the WBcel235 (ce11) *C. elegans* reference genome that were present in the unbackcrossed and backcrossed mutants, but not in the parent strain *acIs101* strain, were identified using in-house scripts. In brief, homozygous variants were called using ‘bcftools’. Any variants that were identified in both the parent *acIs101* strain and the forward genetic mutants were removed using ‘bedtools’. Finally, homozygous variants were annotated using ‘snpEff’ with *C. elegans* reference genome WBcel235.99.

### *C. elegans* strain construction

All CRISPR genome editing was performed as previously described.^66^ CRISPR-Cas9 editing with single-stranded oligodeoxynucleotide (ssODN) homology-directed repair was used to generate *rotr-1(ums53)* and ROTR-1 epitope-tagged strains. All CRISPR reagents were purchased from Integrated DNA Technologies. Target guide sequences were selected using the CHOPCHOP web tool. The ssODN repair templates contained indicated deletions with 35 bp flanking homology arms. ssODN sequences are listed in Table S4. The F1 progeny were screened for Rol phenotypes 3 to 4 days after injection and then for indicated edits using PCR and Sanger sequencing. Primer sequences used for genotyping are listed in Table S4.

Generation of transgenic rescue strains was performed as previously described.^17,19^ For endogenous transgenic rescue of *rotr-1,* the *rotr-1* promoter (∼2kb upstream of the ATG start codon), *rotr-1*, and the 5’ and 3’ untranslated regions (UTRs) were amplified and cloned into a pUC19 vector. The plasmid was then microinjected at 25 ng/µL with 5 ng/µL pCFJ104 (*myo-3*p::*mCherry*::*unc54*) and 120 ng/µL empty pUC19 plasmid. For tissue-specific transgenic rescue strains, the *rotr-1* gene, including both the 5’ and 3’ UTRs were fused to either the *col-10* promoter (for hypodermal rescue), *vha-6* promoter (for intestinal rescue), or the *sng-1* promoter (for pan-neuronal rescue) via Gibson assembly. Plasmids were then microinjected into the *rotr-1(ums53); agIs219* mutants at 10 ng/µL with 5 ng/µL pCFJ104 (*myo-3*p::*mCherry*::*unc-54*) and 135 ng/µL empty pUC19 plasmid.

### Feeding RNAi

*C. elegans* were fed *E. coli* HT115 expressing dsRNA targeting the genes of interest, as previously described with modifications.^51,67,68^ In brief, HT115 bacteria expressing specific dsRNA were grown on LB agar containing 50 µg/mL ampicillin and 15 µg/mL tetracycline at 37 °C overnight. Colonies were then inoculated in LB broth containing 50 µg/mL ampicillin overnight at 37 °C for 16-18 hrs with shaking at 250 rpm. Overnight cultures were then seeded onto NGM plates containing 5 mM IPTG and 50 µg/mL carbenicillin and incubated for 16-18 hrs at 37 °C. Synchronized L1 animals were then transferred onto NGM plates with the grown bacteria and allowed to mature to the L4 stage.

For Figures 3G and 3H, *C. elegans* animals were treated with two-generation RNAi. Animals were first allowed to mature from L1 to gravid adults on individual RNAi clones and then treated with hypochlorite to release eggs. Eggs were allowed to hatch overnight and synchronized L1s were dropped onto the same RNAi clones that their parents were grown on.

### LysoTracker Red assays

Animals were stained with LysoTracker Red (Thermo Fisher) as previously described.^27,28^ To stain animals with LysoTracker Red, 60 mm NGM plates (about 10 mL media) were first seeded with *E. coli* OP50 or HT115 and allowed to dry. Stocks were then diluted 1:10 in M9W (100 µM). 100 µL of each dye was then added on top of the dried bacteria and the dye was allowed to percolate through the plates for 1-2 hrs (final concentration 1 µM). L1 synchronized animals were then dropped on NGM plates containing either 1-2% DMSO or 1 µM LysoTracker Red and allowed to grow in the dark until the L4 stage. About 1-2 hrs before imaging, animals were transferred to NGM plates containing freshly seeded *E. coli* OP50. Stained animals were visualized using the Zeiss AXIO Imager Z2 microscope with a Zeiss Axiocam 506 mono camera and Zen 2.5 (Zeiss) software.

### Microscopy and image analysis

For GFP fluorescence imaging, nematodes were mounted onto 2% agarose pads, paralyzed with 50 mM tetramisole (Sigma), and imaged using a Zeiss AXIO Imager Z2 microscope with a Zeiss Axiocam 506 mono camera and Zen 2.5 (Zeiss) software. For TIR-1::wrmScarlet imaging in Figure 1D, animals were mounted onto 2% agarose pads, paralyzed with 50 mM tetramisole (Sigma), and sealed with coverslips and VALAP. Slides were then inverted and imaged on the THUNDER Imager (Leica DMi8, inverted microscope) with a Leica 63X objective and LasX software (Leica). For Figure 3E, animals were imaged using the Zeiss AXIO Imager Z2 microscope with a Zeiss Axiocam 506 mono camera and Zen 2.5 (Zeiss) software.

TIR-1::wrmScarlet puncta quantification was performed as described previously.^12^

For Figures 6C-F, animals were imaged on the THUNDER Imager (Leica) and images were deconvoluted with Thunder. The diameter of PGP-2::GFP and LMP-1::GFP vesicles were then quantified on ImageJ (Fiji). For each animal, about 10 GFP+ vesicles were randomly identified in the last intestinal cell pair. The vesicular diameters were determined by using the line tool across the widest part of each vesicle. The measured diameters for each animal were averaged and displayed as a single datapoint. Source data can be found in Table S3.

### Gene expression analyses and bioinformatics

RNA-sequencing and data analysis were performed as previously described.^12,69^ Briefly, synchronized wild-type T24B8.5p::*gfp* or *rotr-1(ums53);*T24B8.5p::*gfp* L1 stage *C. elegans* were grown to the L4 stage on NGM plates seeded with *E. coli* OP50. At the L4 stage, animals were then washed to “slow-kill” plates containing *E. coli* OP50 or *P. aeruginosa* PA14. Animals were exposed to each condition for 4 hrs at 25 °C. Animals were then harvested by washing with M9 multiple times before RNA was isolated using TriReagent (Sigma-Aldrich), column purified (Qiagen), and analyzed by 100 bp paired-end mRNA-sequencing using the BGISEQ-500 platform (BGIAmericasCorp) with >20 million reads per sample. The quality of raw sequencing data was evaluated by FastQC (version 0.11.5), and clean reads were aligned to the *C. elegans* reference genome (WBcel235) and quantified using Kallisto (version 0.45.0)^57^. Differentially expressed genes were identified using Sleuth (version 0.30.0)^58^. Pearson correlation statistical analysis was performed using Prism 10. Gene set enrichment analysis of RNA-seq was performed using WormCat^60^ for annotation of *C. elegans* gene categories and GSEA (version 4.2.3)^59^. For GSEA, each differentially regulated gene was given a *π*-value.^49^ The *π*-value was calculated as:

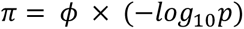

where *ϕ* equals log_2_(fold-change) and *p* equals the *P*-value. Genes were then preranked from the highest *π* value to the lowest *π* value.

For qRT-PCR studies, RNA was extracted from about 3000 L4 animals and reverse transcribed to cDNA using the iScript cDNA Synthesis Kit (Bio-Rad). cDNA was then analyzed on the CFX384 thermocycler (Bio-Rad) with primers described in Table S4. All values were normalized to the geometric mean of the housekeeping control genes *act-3* and *snb-1*. Relative expression was then calculated using the Pfaffl method.^70^

### Immunoblot analyses

Protein lysates were prepared using a Teflon Dounce homogenizer from 5-10,000 *C. elegans* animals grown to the L4 larval stage on NGM plates seeded with *E. coli* OP50, as previously described.^12,69^ LDS Sample Buffer (Thermo Fisher Scientific) was added to a concentration of 1X with 5% β-mercaptoethanol. All samples were incubated at 70 °C for 10 minutes. Total protein from each sample was resolved on NuPage Bis-Tris 4–12% gels (Invitrogen), for detection of phosphorylated p38 PMK-1, total p38 PMK-1 and ROTR-1, or NuPage Tris-Acetate 3-8% gels (Invitrogen), for detection of TIR-1::3xFLAG. Protein was then transferred to 0.2 µM nitrocellulose membranes (Bio-Rad) and blocked with 5% milk in 1x TBS + 0.2% Tween-20 for one hour. Blots were then probed with a 1:1000 dilution of mouse monoclonal anti-FLAG M2 (Sigma, #F1804), 1:1000 phospho p38 PMK-1 (Cell Signaling Technology, #9211), 1:1000 total PMK-1,^23^ 1:2000 mouse monoclonal anti-alpha-Tubulin (Sigma, #T5168), overnight at 4 °C. A polyclonal antibody against the ROTR-1 protein was raised using the peptide LDRSPPSDDGTQKV (ROTR-1 amino acids 288 to 301) in rabbit (Thermo Fisher Scientific). Anti-ROTR-1 sera was used at a dilution of 1:1000. We confirmed that anti-ROTR-1 sera was specific for ROTR-1 using *rotr-1(ums53)* (Figure S2B). For detection of 3xFLAG-tagged ROTR-1, blots were probed with 1:500 dilution of mouse monoclonal anti-FLAG M2. For detection of 3xHA-tagged ROTR-1, blots were probed with 1:500 anti-HA (Roche, #11867423001). Anti-mouse IgG-HRP (Abcam, #ab6789), anti-rabbit IgG-HRP (Cell Signaling Technology, #7074), or anti-rat IgG-HRP (Abcam, #ab97057) secondary antibodies were used at a dilution of 1:10,000 to detect primary antibodies. Blots were then developed with the addition of SuperSignal™ West Pico or West Femto PLUS Chemiluminescent Substrate (Thermo Fisher Scientific) and visualized using a ChemiDoc MP Imaging System (Bio-Rad). Band intensities were quantified using ImageJ (Fiji).

NativePAGE analysis was performed as previously described. Protein lysates were prepared in 1x NativePAGE buffer (Invitrogen), 1% digitonin, and HALT protease inhibitor (Thermo Fisher). Coomassie G-250 additive was added to samples and loaded onto NativePAGE 3-12% Bis-Tris gels (Invitrogen). Samples were first run in dark blue cathode buffer (inner chamber) until the front ran 1/3 down the gel. The buffer was then switched to the light blue cathode buffer. Protein was then transferred onto 0.2 µm PVDF membranes overnight at 4°C. After transfer, membranes were placed in 8% acetic acid and fixed for 15 minutes at room temperature on a rocker. The blot was then destained with 50% methanol/10% acetic acid until the membrane was white and blocked with 5% milk in 1x TBS + 0.2% Tween-20 for 1 hr at room temperature on a rocker. Blots were then transferred to mouse anti-FLAG M2 (Sigma, #F1804) antibody overnight at 4°C. The following day, blots were washed and transferred to anti-mouse IgG-HRP (Abcam, #ab6789) for 1 hr at room temperature on a rocker. Blots were then developed with the addition of SuperSignal^TM^ West Femto PLUS Chemiluminescent Substrate (Thermo Fisher Scientific) and visualized using a ChemiDoc MP Imaging System (Bio-Rad).

### Development and lifespan assays

Development assays were performed as previously described.^16,17^ Briefly, around 200 synchronized L1 animals were grown on *E. coli* OP50 for 48 hrs at 20 °C. Larval stages were then visually quantified under a dissecting microscope. Animals older than the L4 stage were identified and the percent L4+ was calculated. Each genotype had four biological replicates. All source data can be found in Table S3.

For lifespan assays, L4 animals from each genotype were transferred onto NGM plates containing 0.1 mg/mL 5-fluorodeoxyuridine (FUDR), to prevent progeny from hatching, and grown at 20 °C. Live animals were scored daily until all animals on each plate died. Any plates with visible contamination were removed from analysis. Sample size (n), mean lifespan and statistics can be found in Table S1.

### *C. elegans* pathogenesis assays

“Slow-killing” *P. aeruginosa* infection experiments were performed as previously described.^71^ In brief, *P. aeruginosa* was grown as described above and 10 µL overnight culture was spread onto the center of 35-mm tissue culture plates containing 4 mL slow-kill agar (0.35% Bacto-peptone, 0.3% sodium chloride, 1.7% agar, 5 µg/mL cholesterol, 25 mM potassium phosphate, 1 mM magnesium sulfate, 1 mM calcium chloride). Plates were then incubated for 24 hrs at 37 °C followed by 24 hrs at 25 °C. *C. elegans* animals at the L4 larval stage were then transferred to *P. aeruginosa* slow-kill plates containing 0.1 mg/mL FUDR. Dead animals were scored twice daily until completion. Three trials of the assay were performed. Sample sizes, mean survival, and p-values for all trials are shown in Table S1.

### Quantification, statistical analysis, and visualization

Differences in the survival of *C. elegans* in the *P. aeruginosa* pathogenesis assays or lifespan assays were determined with the log-rank test after survival curves were estimated for each group with the Kaplan-Meier method. OASIS 2 was used for these statistical analyses.^55^ Statistical hypothesis testing was performed with Prism 10 (GraphPad Software) using methods indicated in the figure legends. Table S3 contains all source data and statistical analysis methods and results. Sample sizes, survival, and p-values for all trials are shown in Table S1.

## SUPPORTING INFORMATION

**Figure S1.**
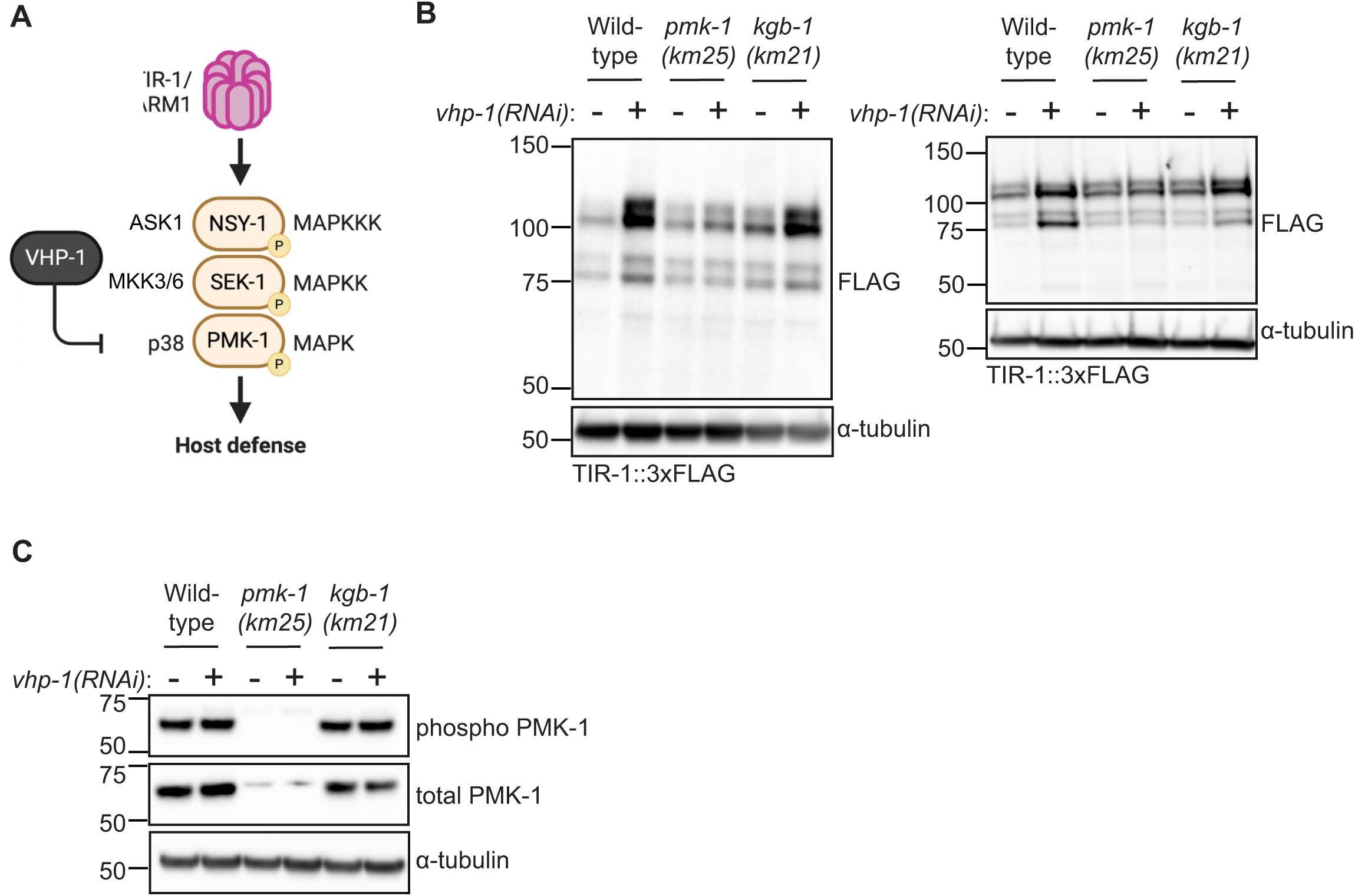
Feedforward activation of the p38 PMK-1 pathway potentiates innate immune defenses by promoting TIR-1/SARM1 multimerization. **(A)** Schematic of the p38 PMK-1 pathway. Mammalian homologs are noted on the left of the main pathway. **(B-C)** Immunoblots of whole cell lysates from wild-type, *pmk-1(km25)*, and *kgb-1(km21)* animals in the TIR-1::3xFLAG background treated with vector control or *vhp-1(RNAi)* and probed for **(B)** anti-FLAG and anti-α-tubulin, or **(C)** anti-phosphorylated PMK-1, anti-total PMK-1 and anti-α-tubulin antibodies. Related to Fig. 1.

**Figure S2.**
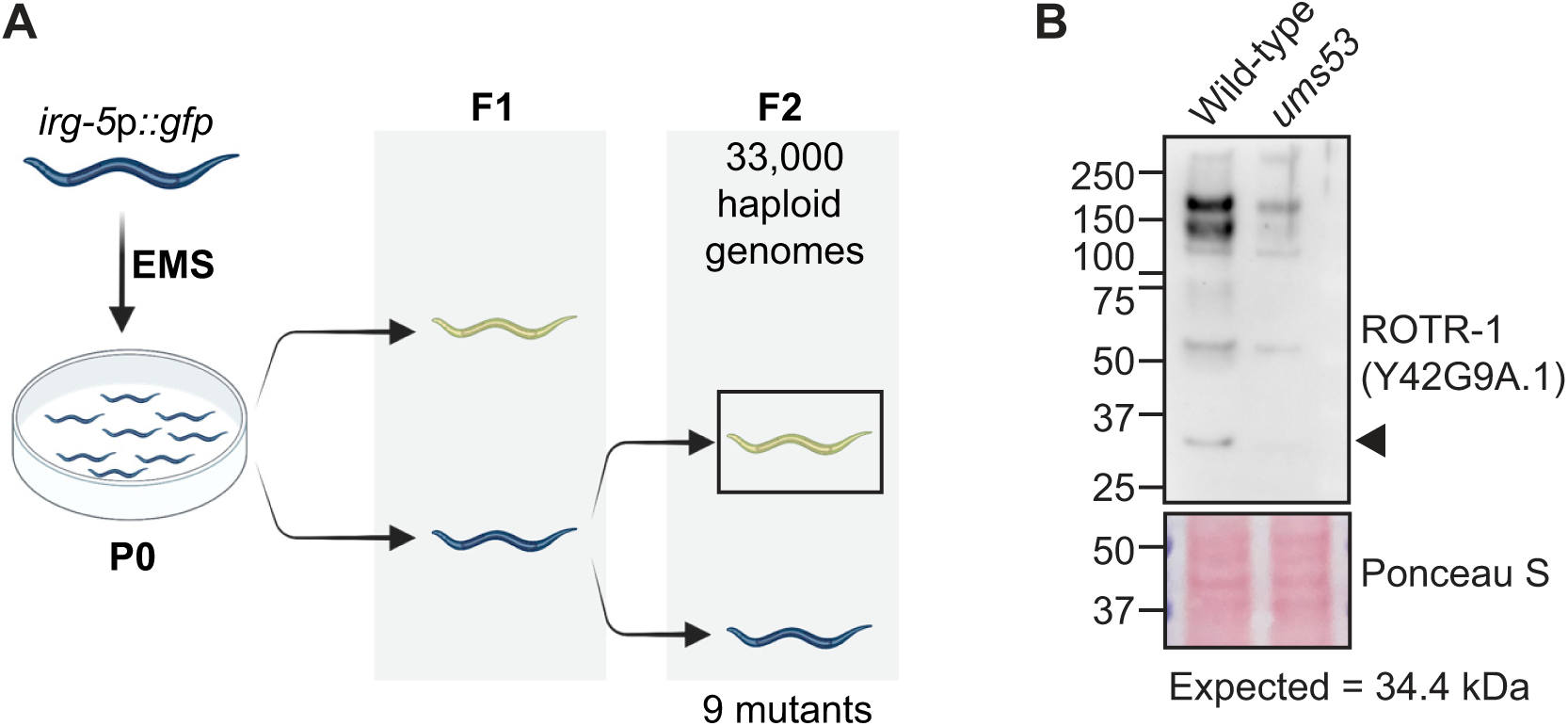
Forward genetic screen identifies *rotr-1*, a novel suppressor of immune gene transcription. **(A)** A schematic of the forward genetic screen using the mutagen ethyl methanesulfonate (EMS). Approximately 33,000 haploid genomes in the F2 generation were screened for constitutive expression of bright GFP fluorescence. **(B)** Immunoblot of wild-type and *rotr-1(ums53)* mutants probed with anti-ROTR-1 sera and stained with Ponceau S. Related to Fig. 2.

**Figure S3.**
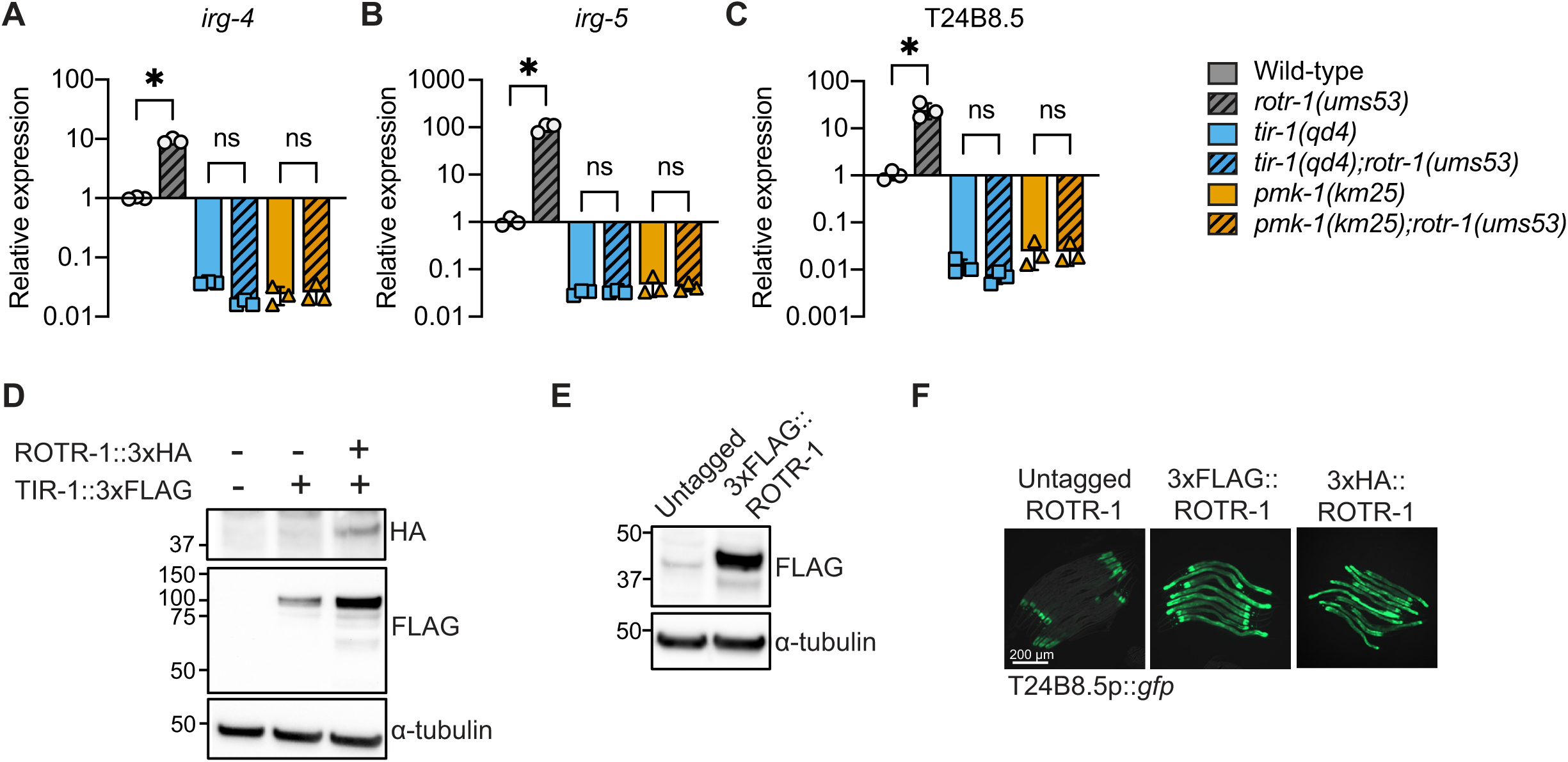
ROTR-1 suppresses feedforward activation of p38 PMK-1 innate immunity. **(A-C)** qRT-PCR analysis of *irg-4* **(A)**, *irg-5* **(B)**, and T24B8.5 **(C)** in indicated genotypes. *equals p<0.05 (one-way ANOVA with Tukey’s multiple comparisons test). ns=not significant. **(D)** Immunoblot of whole cell lysates from wild-type untagged, TIR-1::3xFLAG-tagged, and C-terminus-tagged ROTR-1::3xHA in TIR-1::3xFLAG-tagged animals probed with anti-HA, anti-FLAG, and anti-α-tubulin antibodies. **(E)** Immunoblot of wild-type untagged and N-terminus-tagged 3xFLAG::ROTR-1 probed with anti-FLAG and anti-α-tubulin antibodies. **(F)** Images of T24B8.5p::*gfp* immune reporter expression of untagged and N-terminus-tagged 3xFLAG::ROTR-1 and 3xHA::ROTR-1 animals. Related to Fig. 3.

**Figure S4.**
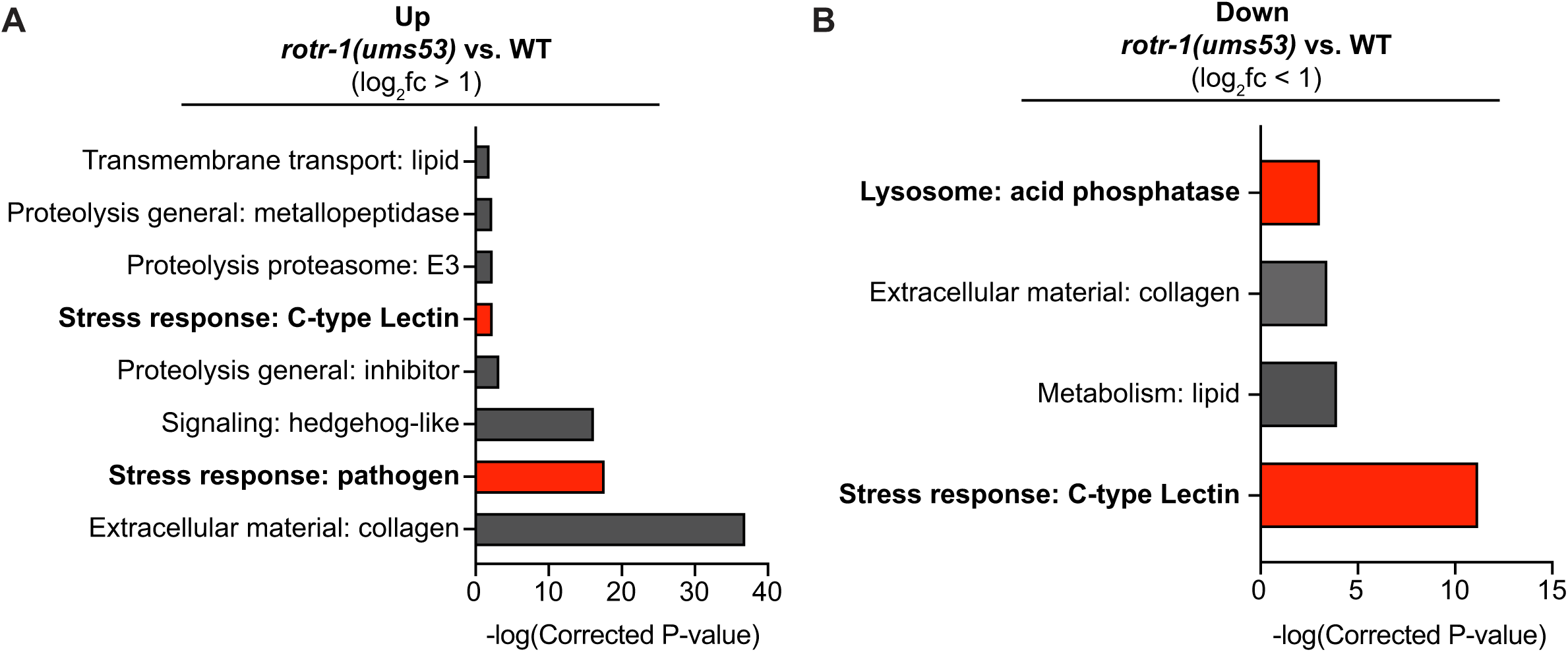
ROTR-1 supports the integrity of lysosome-related organelles, which restrains feedforward activation of p38 PMK-1 innate immunity. **(A-B)** WormCat analyses of significantly differentially expressed upregulated (**A)** or downregulated **(B)** genes in *rotr-1(ums53)* mutants compared to wild-type animals. Bar size represents -log(corrected P-value). Bars of interest are highlights in red. See also Table S3. Related to Fig. 6.

Table S1A-B. Sample sizes, survival, and p values for (A) *C. elegans* pathogenesis assays and (B) *C. elegans* lifespan assays.

Table S2. Differentially expressed genes from RNA-sequencing experiments.

Table S3. Source data and statistical tests used for each figure and supplemental figure.

Table S4. Primer, crRNA guide, and ssODN sequences designed for this study.

